# Markov State Models and NMR Uncover an Overlooked Allosteric Loop in p53

**DOI:** 10.1101/2020.10.19.346023

**Authors:** Emilia P. Barros, Özlem Demir, Jenaro Soto, Melanie J. Cocco, Rommie E. Amaro

## Abstract

The tumor suppressor p53 is the most frequently mutated gene in human cancer, and thus reactivation of mutated p53 is a promising avenue for cancer therapy. Analysis of wildtype p53 and the Y220C cancer mutant long-timescale molecular dynamics simulations with Markov state models and validation by NMR relaxation studies has uncovered the involvement of loop L6 in the slowest motions of the protein. Due to its distant location from the DNA-binding surface, the conformational dynamics of this loop has so far remained largely unexplored. We observe mutation-induced stabilization of alternate L6 conformations, distinct from all experimentally-determined structures, in which the loop is both extended and located further away from the DNA-interacting surface. Additionally, the effect of the L6-adjacent Y220C mutation on the conformational landscape of the functionally-important loop L1 suggests an allosteric role to this dynamic loop and the inactivation mechanism of the mutation. Finally, the simulations reveal a novel Y220C cryptic pocket that can be targeted for p53 rescue efforts. Our approach exemplifies the power of the MSM methodology for uncovering intrinsic dynamic and kinetic differences among distinct protein ensembles, such as for the investigation of mutation effects on protein function.

## INTRODUCTION

The transcription factor p53, known as the “guardian of the genome”, is the most important tumor suppressor in humans due to its regulation of a wide range of cellular activities ^1,2^. Loss of function through p53 missense mutations is associated with progression of about half of human cancers ^3,4^, and reactivation of mutated p53 is emerging as an exciting possibility in cancer treatment as it has been found to lead to tumor regression ^5–9^. More than 90% of the cancer mutations are found in the DNA-binding domain (DBD) of p53 ^10^ (Figure 1a), but the mechanism through which a single mutation affects function is far from resolved. Moreover, the current paradigm is that p53 mutants are not equivalent proteins, but rather have distinct individual profiles in terms of loss of wildtype activity and acquisition of unique tumor-promoting gain of functions ^11,12^. Oncogenic variations can be classified as contact mutations, which lead to loss of function due to disruption of the DNA interaction network ^13^, or structural mutations, which cause perturbations to the DBD and inactivation through destabilization of the protein structure, unfolding and aggregation ^14–17^.

**Figure 1.**
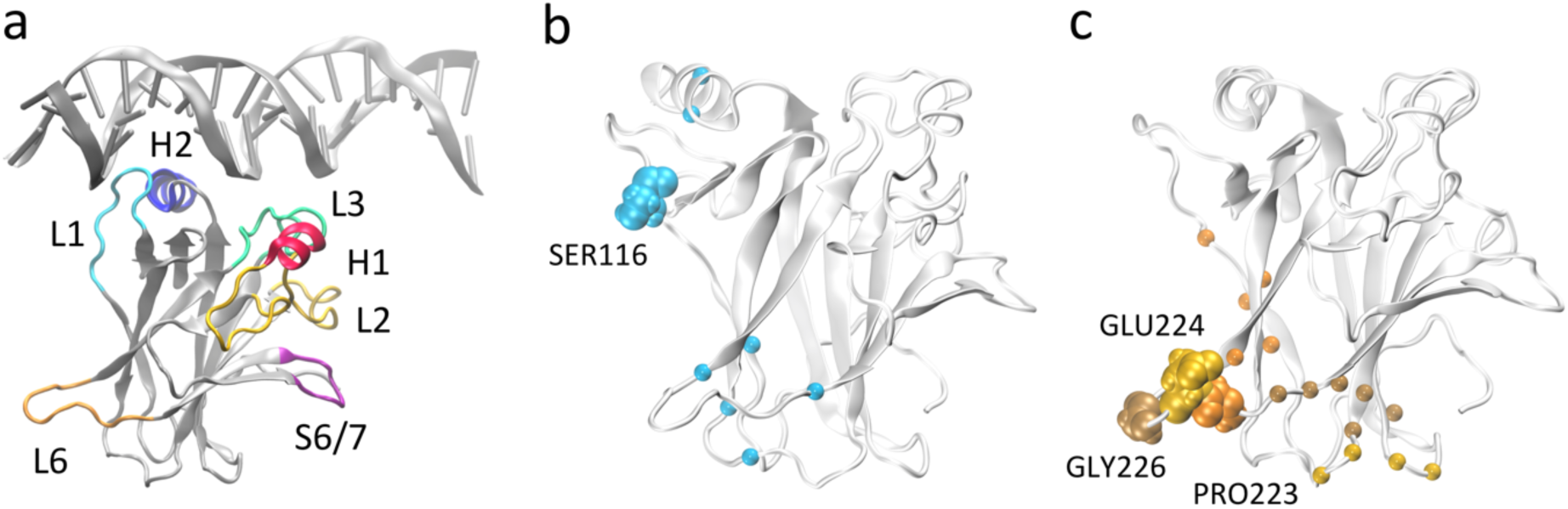
p53 DNA binding domain (a) Monomeric p53 DNA-binding domain in complex with DNA (from PDB 1TSR) with important functional regions highlighted. (b) and (c) Residues used for MSM construction based on pairwise distances, with L1 (b) and L6 (c) anchor residues highlighted in VDW representation. The Cα carbons of the residues that were selected as the second member of the pair with the respective anchor are represented as spheres.

A strategy currently pursued for reactivation of structural mutants is the development of small molecules that bind to the folded state of the protein and restore wildtype p53 conformation and function, with promising results achieved by several groups ^18–27^. Even in proof-of-concept studies, the success of small molecules in reactivating one or a few specific mutants but not others points to the unique behavior of each p53 cancer mutant. In this way, exploring and characterizing the dynamic behavior of different p53 mutants as individual entities promises to open up novel therapeutic opportunities for mutant-specific p53 reactivation.

One such mutant targeted for small molecule reactivation is Y220C, a structural mutant responsible for about 100,000 new cancer cases every year ^14^ and the most frequent p53 cancer mutation observed outside the DNA-binding interface of the protein. The mutation of the bulky tyrosine to the smaller cysteine induces the formation of a crevice in the protein surface that is amenable to small molecule binding ^28–30^, but so far current efforts have failed to yield very high affinity binders ^31–33^.

While the use of molecular dynamics (MD) simulations has allowed the successful identification of druggable pockets on the protein surface of the p53 core domain ^26,32,34^, our understanding of the protein conformational ensemble and dynamics is restricted by sampling limitations. This leaves large regions of the energy landscape unexplored which may include many of the functionally important slower motions. Already, relatively short-scale MD simulations of Y220C have evidenced the flexibility of the protein and the Y220C pocket ^32^. A comprehensive model of p53’s conformational ensemble and the underlying free energy landscape is desirable as it will allow the understanding of the dynamics of key loops and druggable pockets and their role in the overall function and motions of the protein. To help overcome this sampling limitation, we employ here the Markov state model (MSM) methodology in conjunction with extensive MD simulations for the investigation of the conformational dynamics of wildtype p53 and the Y220C mutant.

MSMs allow the integration of multiple MD simulations into a single model of the protein conformational ensemble that contains key thermodynamic and kinetic properties in addition to retaining the atomic level details of the system ^35–38^. Because the MSM is built on the transitions between states, the information from multiple MD simulations of the same system can be combined into a single model and no single simulation has to explore all the states. Importantly, as the equilibrium distribution of states can be derived from the final model, the thermodynamics of the states can be determined, in addition to kinetics, principal motions, and transition pathways of the protein conformational ensemble.

Our computational models, followed by experimental validation with NMR relaxation studies, allow for the first time a thorough exploration of the conformational ensemble of p53 DBD and uncovers the involvement of a loop located away from the DNA binding interface, L6 (residues 221-230), in the slowest dynamics of the wildtype protein. This loop is adjacent to the Y220C mutation, but our models indicate that the mutation affects the conformational landscape of not only L6 but also of the essential DNA-interacting L1 loop. The existence of allosteric communication between the two loops is suggested and provides a mechanistic rationale to the effect of the mutation on the activity of p53. Moreover, analysis of the conformational diversity of L6 evidences the existence of very distinct loop conformations than previously observed experimentally, and the identification of a novel cryptic pocket nestled in the extended conformation of L6 that could be exploited for mutant-specific drug design efforts. Our work emphasizes the ability of MSMs to explore in detail protein conformational landscapes, uncover hidden states inaccessible to experiments and inform on mutation or other environmental effects (such as ligand binding or pH) on protein function.

## RESULTS AND DISCUSSION

### L6 is the slowest loop in p53 DBD dynamics

Markov state models provide a framework for exploring protein dynamics with atomic resolution beyond the timescales typically accessed by molecular dynamics simulations. A crucial step when integrating molecular dynamics trajectories for model building is the selection of features used to discretize the protein conformations sampled, which decreases the dimensionality of the conformational space while still allowing for discrimination between distinct states and appropriate representation of the relevant motions. For a general understanding of the protein conformational ensemble, the task can become challenging due to the conflict between the large degrees of freedom required to describe the protein ensemble and the need to limit the feature dimension to a small, tractable number for model building.

To investigate the basal dynamics of wildtype p53, we employed an unbiased method that started from computing all possible pairwise distances (18,336 features), and iteratively performed time-lagged Independent Components Analysis (tICA) ^47^ to identify the linear combination of features that describe the slowest motions of the system, followed by elimination of the features with low tICA correlation (Supplementary Table 1). Using this methodology we arrived at a final number of 24 pairs (Supplementary Table 2). tICA is useful in the data processing for MSM construction as it maximizes the feature combination to yield kinetically relevant independent components (tICs) representing the slowest degrees of freedom in the system. Despite starting from all possible pairwise distances and including no directed selection of features besides the elimination of pairs that involve the clipped terminal residues or that are consistently too close (< 3Å) or too far (>10 Å) throughout the whole simulations, the final set consisted of interacting pairs centered around loops L1 (residues 113-123) and L6 (residues 221-230). All pairs involved at least one residue located in either L1 (Ser116) or L6 (Pro223, Glu224, Gly226), hereafter referred to as L1 and L6 anchor residues, respectively (Figure 1 b-c, Supplementary Table 2).

The presence of the repeated anchor residues in the final feature pairs suggests that loops L1 and L6 are involved in the slowest and most significant motions of the protein. Loop L1 is known as a dynamic and biologically important motif for p53 function, having been observed experimentally and computationally in two very distinct extended (Figure 1a) and recessed conformations ^59–61^. The identification of the relevance of loop L6, on the other hand, sheds a light on a relatively unexplored region of p53. Not much attention has been given to the role of this structural motif, possibly because of its distance from the DNA-binding surface. However, elevated B factors in p53 crystal structures and NMR NOE values ^58^ point to its intrinsic dynamics, and flexibility in this loop was observed in an early short simulation of wildtype p53, even though implications for functionality were not explored as the motion was deemed to stem from a lack of crystal packing ^62^.

For a comparison of the conformational landscapes of wildtype and Y220C, the conformations explored by each of the simulations and represented by the 24 features were jointly used as input for tICA, and the resulting free energy landscapes are shown in Figure 2a. The tIC independent component space is therefore the same for wildtype and mutant free energy landscapes and allows for a direct comparison of the conformational ensemble explored by each system. The wildtype simulation presents two preferred states, corresponding to the minima in the free energy landscape. The main distinction between them are the conformations of L1 and indicate the same recessed and extended L1 conformations that have been previously observed (Figure 1b). Interestingly, the pairwise features used for construction of the map align with the tICA components in this transformed coordinate space, permitting a direct interpretation in terms of protein conformation: tIC1 is closely correlated with features that include L6 anchors, and tIC2 is more closely correlated with features involving the L1 anchor, Ser116 (Figure 2c), such that tIC1 and 2 are associated with the relative motions of L6 and L1, respectively. Moreover, visual inspection of the conformations distributed on the free energy landscape evidence that smaller values of each of the tICs describe conformations with extended loops, while larger values describe recessed loop conformations (novel conformations are discussed in more detail in following sections).

**Figure 2.**
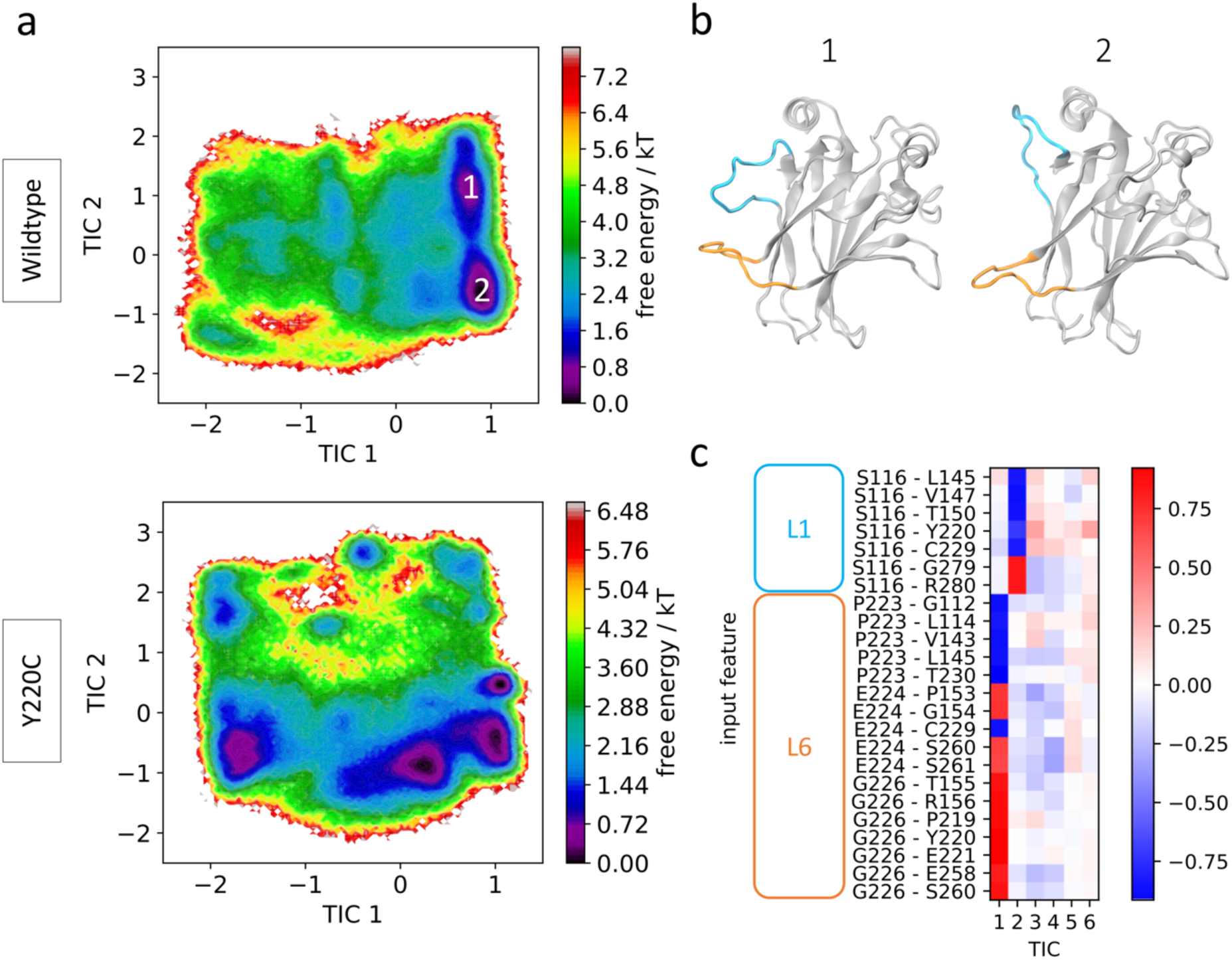
Wildtype and Y220C simulation results. (a) Free energy landscape of wildtype (top) and Y220C (bottom) in terms of tICA components (tICs). (b) Representative conformations from the wildtype preferred states. Loops L1 and L6 are highlighted in blue and orange, respectively. (c) Feature correlation with the first five tICA components. Pairwise distances involving L1 or L6 loop anchor residues are indicated.

Since the tICs are ordered in terms of slowest to fastest motions, the correspondence of L6 anchor features with the first of the components indicates that transitions involving loop L6 are slower than those for loop L1. To further check the importance of these loops in the relevant motions of the protein, we performed additional tICA analysis specifically incorporating other motifs known to play significant roles in p53 function: helices H1 and H2 and loops L2 and L3, which together with L1 make up the DNA interaction surface, and loop S6/7, recently identified as a flexible region in p53 mutants ^63^ (Figure 1a). Even though several of these loops show pronounced flexibility in the simulations as indicated by Ca RMSF values (Supplementary Figure 5), loops L1 and particularly L6 still dominate the slowest transitions (Supplementary Figure 6). This suggests that, while other regions such as loops L2 and S6/7 may be highly flexible as evidenced by their high RMSF values, they display fast dynamics and act as further evidence to the important role of L6 on the slow dynamics of p53.

### Allosteric communication between L1 and L6

In Figure 1a it can be seen that the Y220C mutation affects not only the conformational landscape of loop L6, where it is located, but also of loop L1 (as represented by tIC 2). This loop is essential for p53 activity as it is involved in key interactions with DNA through hydrogen bonds formed by Lys120 and Ser121 ^61^. Wildtype p53 shows important intrinsic L1 flexibility, but the effect of the mutation on loop L1’s dynamics indicates the existence of possible long-range communication between L1 and L6.

To look into this in more detail, we constructed MSMs for the wildtype and mutant system using only the above identified features that include the L1 anchor, Ser116. The conformational landscape in terms of these 7 features, following tICA transformation, is shown in Figure 3a, and includes the coordinates of experimentally-characterized wildtype and Y220C p53 structures for comparison (Supplementary Tables 3 and 4). Coarse-graining of the structures using Hidden Markov state models identifies the presence of 5 metastable states in each case. Two wildtype metastable states, states A and B, are retained in the mutant system with slight changes to their equilibrium populations (Figure 3b-c). State A is the most populated state in both systems, and shows loop L1 in the most extended-like conformations. In wildtype state B, we see a previously-identified 3-10 helix in the L1 loop, absent in the corresponding Y220C state.

**Figure 3.**
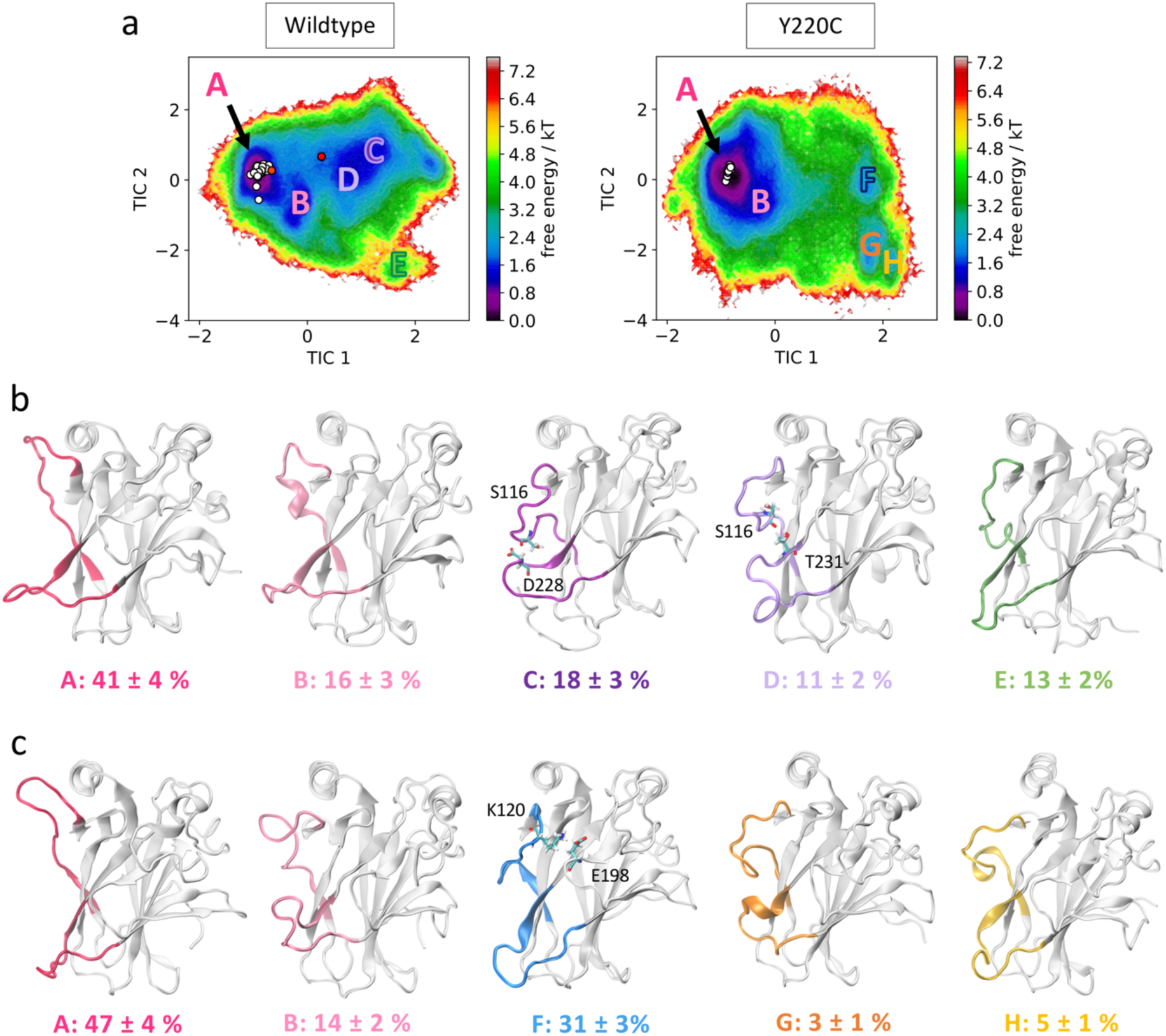
L1-centered MSM (a) Free energy landscape of wildtype (left) and Y220C (right) in terms of the features that describe L1 relative dynamics. Location of metastable states are indicated with letters from A to H. Experimentally resolved DBD structures (X-ray crystallography and NMR) are indicated as white (extended L1 conformation) and red (recessed L1) circles. (b) Conformations from each of the wildtype metastable states. Equilibrium populations and standard deviations are indicated. Residues identified in key interactions are highlighted. (c) Conformations from each of the Y220C metastable states.

The second, shallower wildtype minima, centered at TIC1 = 1, is absent in the Y220C sampled conformations. Indeed, we find that two wildtype metastable states are abrogated by the mutation (states C and D), being substituted by a single state in the Y220C system (state F). These wildtype states show L1 in recessed conformations, and jointly account for 29% of the equilibrium population. Interestingly, in both cases we find that L6 is also organized in a recessed conformation, such that both loops are located in close proximity to each other. Investigation of the loop residues indicates inter-loop hydrogen bonds formed between the side-chain oxygen of Ser116 in L1 and backbone nitrogen of Asp228 (in state C) or side-chain oxygen of Thr231 (state D) in L6 (Figure 3b and Supplementary Figure 7).

Loop L1 in the corresponding Y220C state F, on the other hand, is found to be more collapsed into the protein surface, in a conformation that does not allow for interaction with L6. Rather, a salt bridge between Lys120 in L1 and Glu198 in loop S5/S6 seems to promote the stabilization of this alternate conformation, which accounts for 31% of the Y220C equilibrium population and is the second most populated Y220C state (Figure 3c). The sequestering of the DNA-interacting Lys120 in this significant metastable state could provide a mechanistic explanation to the p53 inactivation effect of the mutation. Furthermore, the conformation-dependent interaction between L1 and L6 identified here suggests the existence of an allosteric communication between them in functional p53, which is disrupted by the mutation.

Finally, we observe a slight destabilization of states located at low values of TIC 2 in the Y220C system, which display loop L1 in extremely-recessed conformations (equilibrium population of 13% for wildtype state E and 8% for Y220C states G and H). There are no persistent L1-L6 interactions in these states. A helical content in loop L6 of Y220C state G seems to be promoted by an inter-L6 hydrogen bond between Ser227 and Thr231.

### Dynamics and druggability of loop L6

The significance of L6 dynamics suggested by the tICA analysis and its effect on the L1 conformational ensemble of wildtype and Y220C prompted us to consider its conformational plasticity in more detail. Figure 4a shows the free energy landscape of the wildtype and Y220C systems now in terms of the tICA components calculated from the 17 L6 anchor features. Again for comparison, we overlay the corresponding coordinates of the X-ray and NMR structures of wildtype and Y220C p53. It is striking how all the previously identified structures are confined to a small area of the map, and the simulations suggest the existence of novel significant protein conformations that remain unexplored to date and could be potentially targeted for drug discovery.

**Figure 4.**
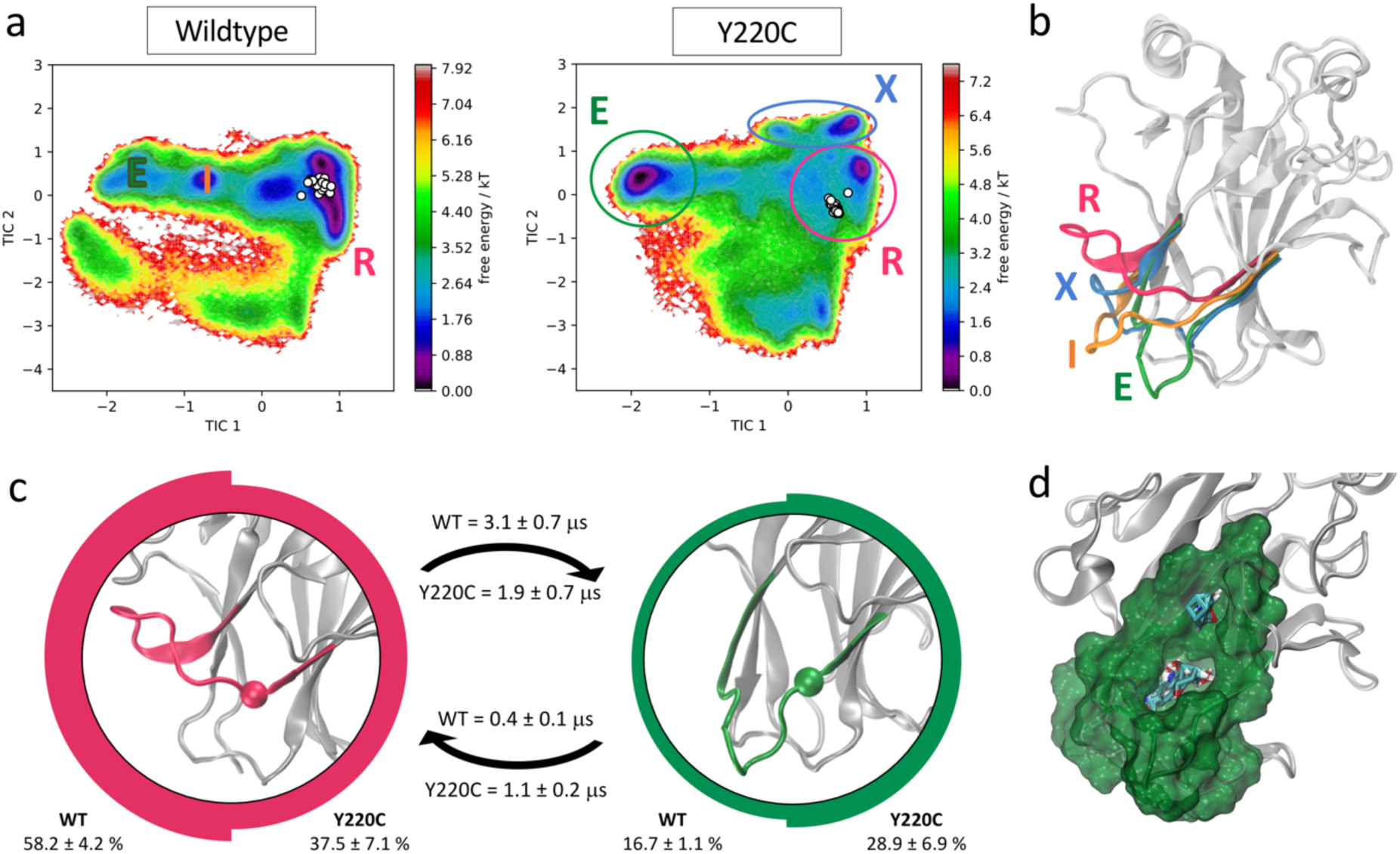
L6-centered MSM. (a) Free energy landscape of wildtype and Y220C systems in terms of L6 features. Experimentally resolved DBD structures (X-ray crystallography and NMR) are indicated as white circles. Populated metastable states are identified. (b) Structural representation of the metastable states identified in the free energy landscapes: recessed (R, pink), intermediate (I, orange), extended (E, green) and mutant exclusive (X, blue) conformations. (c) Equilibrium population and mean first passage times (MFPT) for the two major wildtype and Y220C metastable states. The images at the center of the circles represent the respective state L6 conformation, with the a carbon of the mutated residue highlighted. Thickness of the circle edge is proportional to the equilibrium population in the respective system (wildtype on the left, Y220C on the right). MFPTs of the transitions are indicated above (for wildtype) and below (for Y220C) the respective arrows. (d) Representation of the novel pocket found in the L6 extended conformation and solvent mapping results.

While the experimental structures align with the wildtype low energy well, the mutation leads to the stabilization of multiple alternative loop L6 conformations, including one mutant-exclusive well at high values of tIC1 that is distinct from the experimentally characterized structures. Five metastable states can be identified from the Markov state models for each of the wildtype and Y220C systems. Two populated wildtype metastable states at equilibrium remain significant states in the Y220C ensemble, albeit with changes to their relative equilibrium population and rates of transitions. Three wildtype states are abrogated by the mutation, while we observe the formation of two Y220C-exclusive metastable states. The conformational differences between the highly populated states and potential for drug discovery are explored in more detail below.

#### The mutation induces stabilization of extended L6 conformations

The most populated metastable state in the wildtype ensemble, accounting for over 58% of the population at equilibrium, corresponds to L6 in a recessed conformation similar to that observed experimentally (Figure 4b-c, pink R state). This organization of the loop allows for the formation of a crevice in between loops L6 and S3/S4 upon the substitution of the bulky tyrosine for the much smaller cysteine residue in the Y220C mutant, which results in the pocket currently targeted for p53 rescue ^28–33^.

In several of the mutant frames belonging to this metastable state we observed the opening of a transient channel through loop L6, connecting the crevice to another area of the protein surface. This cryptic pocket has been identified previously by Fersht and co-workers using molecular dynamics simulations ^32^, and in agreement with their studies, we find it to exhibit promising druggable characteristics (as suggested by FTMap ^49^ solvent mapping analysis, Supplementary Figure 8). Exploitation of this channel by small molecules could improve the potency of rescue drugs and increase specificity towards mutant p53, as the channel is unavailable in the wildtype simulations due to the larger volume occupied by the tyrosine residue.

Besides this well-characterized state, the MSMs indicate an additional common metastable state in the wildtype and Y220C ensemble at equilibrium. This metastable state, corresponding to 16.7% of the wildtype population and 28.9% of the Y220C ensemble, exhibits loop L6 in a dramatically different extended conformation (green E state in Figure 4b-c). In this conformation, the crevice underneath L6 typically targeted by small molecules for Y220C reactivation is disrupted. However, visual inspection identified the formation of another cryptic pocket nestled within this loop, promoted by the extended conformation of L6 (Figure 4d). Similar to the mutant-induced crevice, this pocket is only evident in the Y220C simulations due to the presence of the less bulky cysteine in its center. The entrance of the cavity in this case faces the opposite side of the loop relative to the known Y220C crevice, in the direction of the DNA binding surface, and the cavity corresponds to a relatively deep hydrophobic pocket with average volume of 333.5 ± 57.6 Å^3^ with opportunities for hydrogen bonding interaction, as well as other polar interactions in the more solvent-exposed region above L6.

Several hydrogen bonds between loops L6 and S3/S4 (residues 146-155) are found to be established for longer fractions of the simulation in the mutant state, with increases of up to 100x in persistence time, and suggest possible interactions promoting the extended conformation (Supplementary Table 5). Further indication of the stabilization of the extended conformation promoted by the mutation is given by the calculation of mean first passage times (MFPT) between these metastable states: The mutation decreases the mean first passage time from the recessed to the extended L6 conformation by a factor of 1.6, resulting in a faster transition in the Y220C mutant compared to the wildtype, while the MFPT out of the extended conformation and into the recessed increases by more than 2 in the Y220C mutant (Figure 4c).

Finally, the third significantly-populated state in the wildtype ensemble, with a stationary population of 18.3% and corresponding to an intermediate state between the extended and recessed conformations (Figure 4b) is completely abrogated in the Y220C ensemble, such that the recessed-extended transition occurs without an intermediate state for the mutant.

#### Characterization of mutant-exclusive metastable states

The long-timescale exploration of the Y220C mutant dynamics evidenced the existence of two mutant-exclusive states (Figure 4a). Jointly, these metastable states account for 24.7% of the relative Y220C ensemble population, a significant portion of the conformational ensemble that opens up promising avenues for mutant-specific therapeutic opportunities. In these states the loop L6 shows a similar extended conformation to the novel metastable state E described above, but with a “sideways” bend likely promoted by a Thr149-Pro222 interaction (Figure 4b, Supplementary Figure 9a). This bend slightly disrupts the cryptic pocket identified in the fully extended L6 conformation, resulting in a smaller and shallower cavity, but also leads to the formation of a channel across loop L6 and underneath the mutation which reaches across to the protein surface at a different site (Supplementary Figure 9b). Transitions into or out of these states constitute the slowest process in the Y220C MSM, with a timescale of approximately 7.3 ± 2.8 μs.

Taken together, our models suggest a molecular explanation for the reactivation of the Y220C mutant achieved by small molecules ^29–33^: since in the mutant the recessed L6 conformation is destabilized (37.5% of the Y220C population versus 58.2% for wildtype) with a preference for the novel E and X extended conformations, binding of a small molecule into the crevice underneath L6 should prevent the transition towards these extended conformations and could lead to a shift in the equilibrium towards a wildtype-like, recessed loop conformational ensemble. Additionally, as the investigation of the full p53 conformational flexibility suggests a high degree of correlation between L1 and L6 dynamics (Figures 2 and 3), this could further indicate a previously uncharacterized functional link between L6 conformation and p53 rescue.

### NMR relaxation analysis

As an external validation of the loop dynamics identified by the MSMs, we performed NMR relaxation studies to determine flexibility of backbone atoms of the wildtype protein. Measurement of ^15^N NOE, longitudinal (R1), and transverse (R2) relaxation rates were used to obtain generalized order parameters (S^2^). The list of relaxation rates and NOEs are found in Supplementary Table 6. Relaxation rates for some residues could not be obtained due to rapid signal decay (not enough points to fit) or significant signal overlap. We used the program *Modelfree* to determine backbone flexibility based on heteronuclear NOE, R1, R2 measurements ^54,55^. Using the *quadric_diffusion* program ^57^ we found that the best fitted diffusion tensor model was an axial symmetry model. The rotational correlation time (τ_m_) was calculated to be 14.7 ns with an axially symmetric tensor (D_∥/⊥_) = 0.29.

Calculated order parameters show that the most flexible regions are in the loop regions (Figure 5a), and that L6 is the most flexible loop. A similar trend in wildtype backbone order parameters has been previously observed ^58^, although our values are larger in magnitude, possibly due to differences in magnetic field strength in which the experiments were performed. Additionally, the R2/R1 ratio, which provides a qualitative indication of the timescales of motions involving backbone residues, indicates that L6 dynamics contain a slow-motion component that occurs at longer timescales (μs-ms) than that of L1 (ps-ns) (Figure 5b), in agreement with our findings.

**Figure 5.**
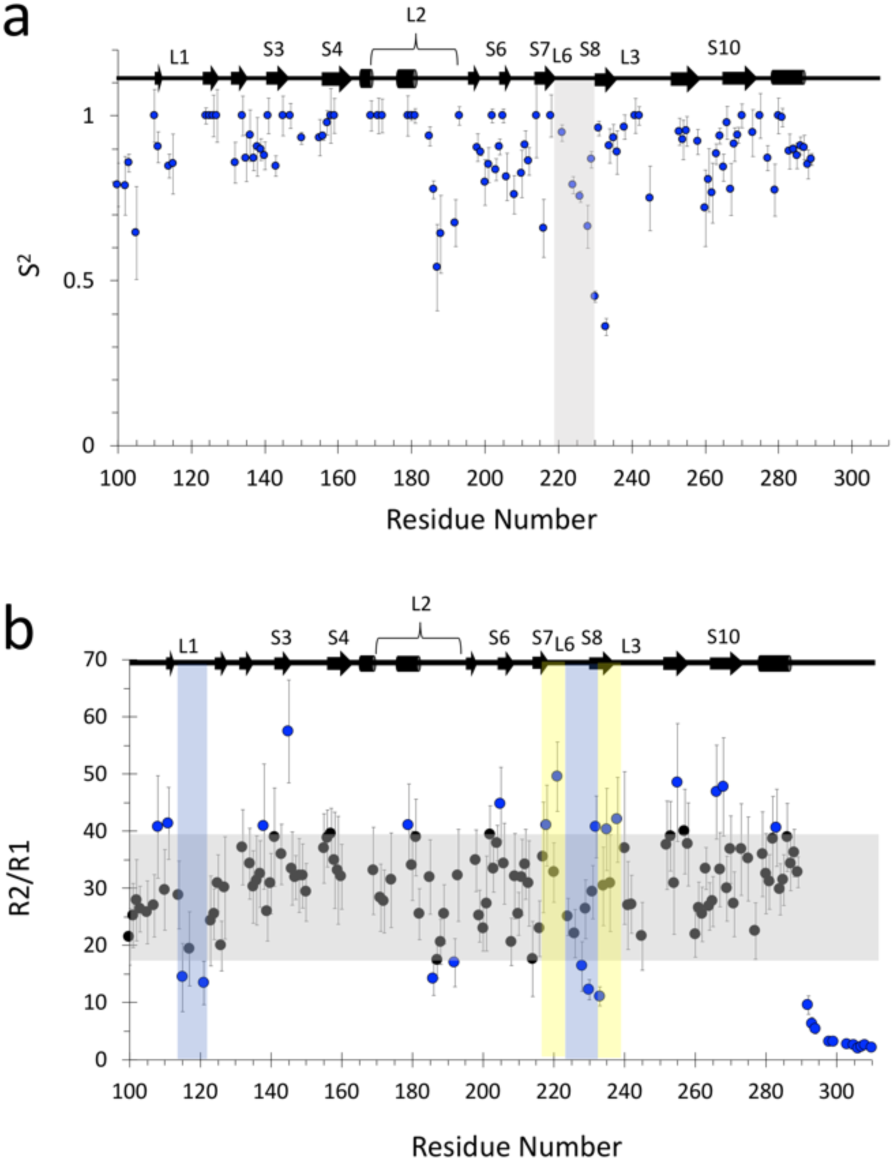
(a) Quantitative characterization of fast dynamics of wildtype p53 DBD. Generalized order parameter (S^2^) for wildtype obtained via *Modelfree* analysis ^54,55^ of relaxation rates R1, R2, and ^1^H-^15^N NOEs at 800 MHz. Vertical boxed area highlights L6. (b) R2/R1 plot for qualitative analysis of backbone dynamics. Grey rectangles highlight mean values ± 1 SD of all R2/. Blue data points highlight the residues that are outside the mean ± 1 SD. Yellow vertical shaded rectangle highlights regions of interest of slow dynamics (μs-ms). Similarly, blue shaded rectangle highlights regions of interest of fast dynamics (ps-ns).

## CONCLUSIONS

Our combined tICA and MSM approach, validated by NMR relaxation measurements, highlights a functional role to the dynamic loop L6, which exhibits motions at longer timescales than other characterized structural motifs and presents potential for mutant-rescue therapeutic opportunities. The conformational landscape suggests some degree of allostery between L6 and the functionally-important loop L1, likely promoted by hydrogen bonds formed when both loops are in the recessed conformation and thus in close proximity to each other.

The Y220C mutation, which characterizes one of the most common cancer mutants, is located at the N terminus of L6, and we find that the mutation promotes the stabilization of novel protein conformations which exhibit loop L6 in extended states instead of the only currently characterized and targeted recessed conformation. The stabilization of the extended conformation induces the formation of a deep hydrophobic pocket within L6 due to the removal of the bulky tyrosine, as well as the population of two mutant-exclusive L6 states that could be explored for mutant-specific therapies.

In summary, the comparison of the dynamics of wildtype and mutant p53 DBD’s using MD simulations and Markov state models evidenced for the first time the existence of functionally-relevant motions involving loop L6 and presents applications for mutant-specific rescue efforts. We anticipate that this approach will be useful in the study of the conformational ensembles of other p53 cancer mutants or protein targets, as a way to provide atomic-level information on these proteins’ motions combined with thermodynamic and kinetic details in tandem with experimental observations.

## MATERIALS AND METHODS

### System set up

The DNA binding domain initial coordinates were taken from chain B of PDB 1TSR, which include p53 amino acids 96 – 289. For the mutant simulations, the tyrosine in position 220 (125 in the clipped domain) was mutated to a cysteine using tleap module in Amber14.^1^ The crystallographic water molecules were retained and each system was solvated in an 8Å TIP3P water box.^2^ The zinc ion and its coordinating residues were modeled using the cationic dummy atom model.^3^ Each system was brought to 0.12 M salt concentration by adding K^+^ and Cl^-^ atoms. The structure file of each system consisted of about 27,220 atoms, which were prepared using Amber FF14SB force field.^1,4^

### Molecular dynamics simulations

The solvated proteins were minimized and equilibrated as described in Malmstrom *et al*,^5^ using the GPU version of Amber 14. To increase the conformational sampling, a round of accelerated MD simulations (aMD)^6^ was performed from the equilibrated structure using Amber14. Each system was simulated for 100 ns and 10 structures were selected for each system by clustering the conformations based on RMSD of the center of mass of each residue using a k-means algorithm in MSMBuilder2^7^ and using the cluster centroids. These 10 structures were used as seeds for short unbiased MD simulations, each performed in triplicate with new starting velocities. After each round of simulation, the joint trajectories were processed for MSM model construction, and new starting coordinates were selected, prioritizing the exploration of new areas in the conformational space, until converged models were obtained based on MSM validation metrics (see below). Individual simulations ranged from 10 to 300 ns in length. In total, the wildtype system was simulated for 89 μs, while Y220C required 63 μs for appropriate model construction.

### Markov state model construction

Simulation data was processed and models were built using PyEMMA^8^, version 3.5.6. Features consisted of pairwise distances, with pairs being selected after a tICA-based iterative process that eliminated redundant pairs located consistently close (< 3Å) or far (>10Å) in all frames of the simulations, as well as pairs involving residues located close to the clipped termini, with low variance (<0.05 Å) and those that accounted for low correlation with the first tICs (Supplementary Table 1). The final feature set consisted of 24 pairs (Supplementary Table 2). Time-independent component analysis (tICA)^9^ was used to process the joint wildtype and Y220C featurized data. Distinct loop-centered Markov state models were constructed using the 17 features that are centered in L6 and the 7 for L1. Discretization was performed with k-means clustering, k = 200, for each system (wildtype and Y220C) separately, and accuracy of the models verified by implied timescale (ITS) plots and Chapman-Kolmogorov tests (Supplementary Figures 6 and 7). The L6 and L1-focused models were constructed with MSM lag time of 10 ns each. The MSMs were then coarse-grained using hidden Markov state models (HMMs) with a lag time of 3 ns (L1) or 2.5 ns (L6), again validated by ITS plots and Chapman-Kolmogorov tests (Supplementary Figures 8 and 9). Standard deviations were calculated using Bayesian hidden Markov state models corresponding to the respective HMMs.

### Pocket characterization

Pocket volume measurements were performed with POVME, version 2.0,^10^ and druggability assessments were based on computational solvent mapping of randomly selected conformations from the MSM metastable states using FTMap.^11^ Existence of hydrogen bonds across the simulations was probed using MDTraj,^12^ with a hydrogen bond defined as established if donor-acceptor distance < 2.5 Å and angle > 120°.

### Conformational landscape comparison with experimental structures

For comparison of the conformations sampled by the simulations with experimentally-resolved p53 structures, we transformed the coordinates of wildtype and Y220C structures solved by X-ray crystallography and NMR spectroscopy applying the same feature and tICA transformation steps used for the L1 and L6 MSMs. A total of 58 wildtype and 54 Y220C chains were selected from 16 and 29 PDB entries, respectively, and are listed in Supplementary Tables 3 and 4. These structures were selected out of all wildtype and Y220C deposited structures as they contained the same number of Cα atoms to the simulation structures and thus could be directly compared in tICA space.

### NMR data collection and sample preparation

WT and mutant p53 constructs were provided by Rainer Brachman. ^15^N-labeled p53 DBDs were prepared similarly to Wong *et al*.^13^ The DBD core domain of human p53 (94-312) and rescue mutants were transformed into BL21 *E. Coli* cells. Bacteria were grown at 30 °C in ^15^N-enriched Neidharts minimal media to a density of 0.8-1.0 OD_(550 nm)_. The temperature was lowered to 18 °C and both IPTG and ZnCl2 were added to a final concentration of 1 mM. Cells were allowed to grow for an additional 6-8 hours and then harvested by centrifugation. The frozen cell pellet was resuspended in 20mM sodium phosphate, pH7.2, 10 mM BME, 0.5 mM PMSF, and lysed using sonication. Cell debris were removed by centrifugation at 4 °C for 30 minutes. The supernatant was applied to a SP-sepharose column and the protein eluted with a gradient of 100 mM-600mM NaCl. Samples were dialyzed into 15 mM potassium chloride, 25 mM sodium phosphate, pH 7.1, 10 mM BME and concentrated to a protein concentration of 400 μM.

All NMR experiments were performed on Varian Inova 800 MHz at 20 °C. NMR data were processed using nmrPipe.^14^ Residues assignments and rate measurements (using peak volumes) in all 2D HSQC spectra were accomplished using CcpNmr Analysis.^15^

3D ^15^N-TOCSY-HSQC (t_m_ = 75 ms) and NOESY-HSQC (t_m_ = 100 ms) were analyzed for assignment of backbone amide resonances. Published WT assignments^13^ were confirmed in spectra of our WT sample.

### Relaxation analysis

T1 and T2 spectra were recorded with relaxation delays of 10, 50, 100, 200, 400, 750, 1000, 1500 ms and 10, 30, 50, 70, 90, 110, 130, 150 ms, respectively. The NOE spectra were recorded with a 5 second irradiation and 3 second delay. R1 and R2 rates were obtain by measuring peak volume as a function of delay time and fitting them to a single exponential function with CcpNmr software.^15^ Errors in relaxations rates were obtain through the covariance method in CcpNmr. R2/R1 error bars are a results of error propagation of both R1 and R2:

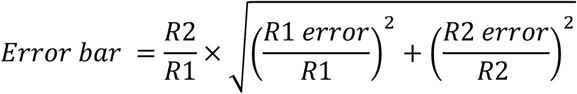

NOE errors were calculated by the following equation:

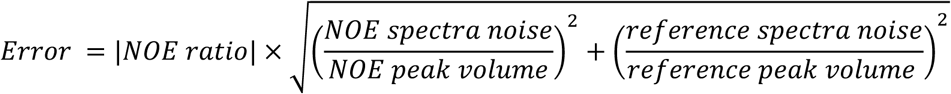

NMR generalized order parameters (S^2^) were obtained by using *Modelfree 4*.*15*^16,17^ in combination with *FASTModelfree*.^18^ Initial estimation of *Modelfree* parameters were obtained through the programs *pdbinertia, r2r1_tm, quadric_diffusion* from the CoMD/NMR website.^19^ Residues with NOE and X^2^ values less than 0.6 and 10, respectively, were excluded from *quadric_diffusion* calculation. Our analysis was performed using the Protein Data Bank (PDB) file 2FEJ. All *FASTModelfree* analysis used an NH bond length of 1.015 Å and chemical shift entropy (CSA) value of −179 ppm similar to the previously published analysis of p53 DBD.^20^ *FASTModelfree* analysis proceeded until all parameters converged.

## ACKNOWLEDGEMENTS

We thank Robert Malmstrom and Nathan Hensley for helpful discussions and assistance with clustering, and Bryn Taylor and Frank Noe for helpful discussions regarding MSM construction. This work was supported by 1R01GM132826-0,1 and funded in part by the National Biomedical Computation Resource (NBCR) through NIH P41 GM103426. J.S. acknowledges training grant NIH-IMSD GM055246 for support.

## Supplementary Information

**Supplementary Table 1.**
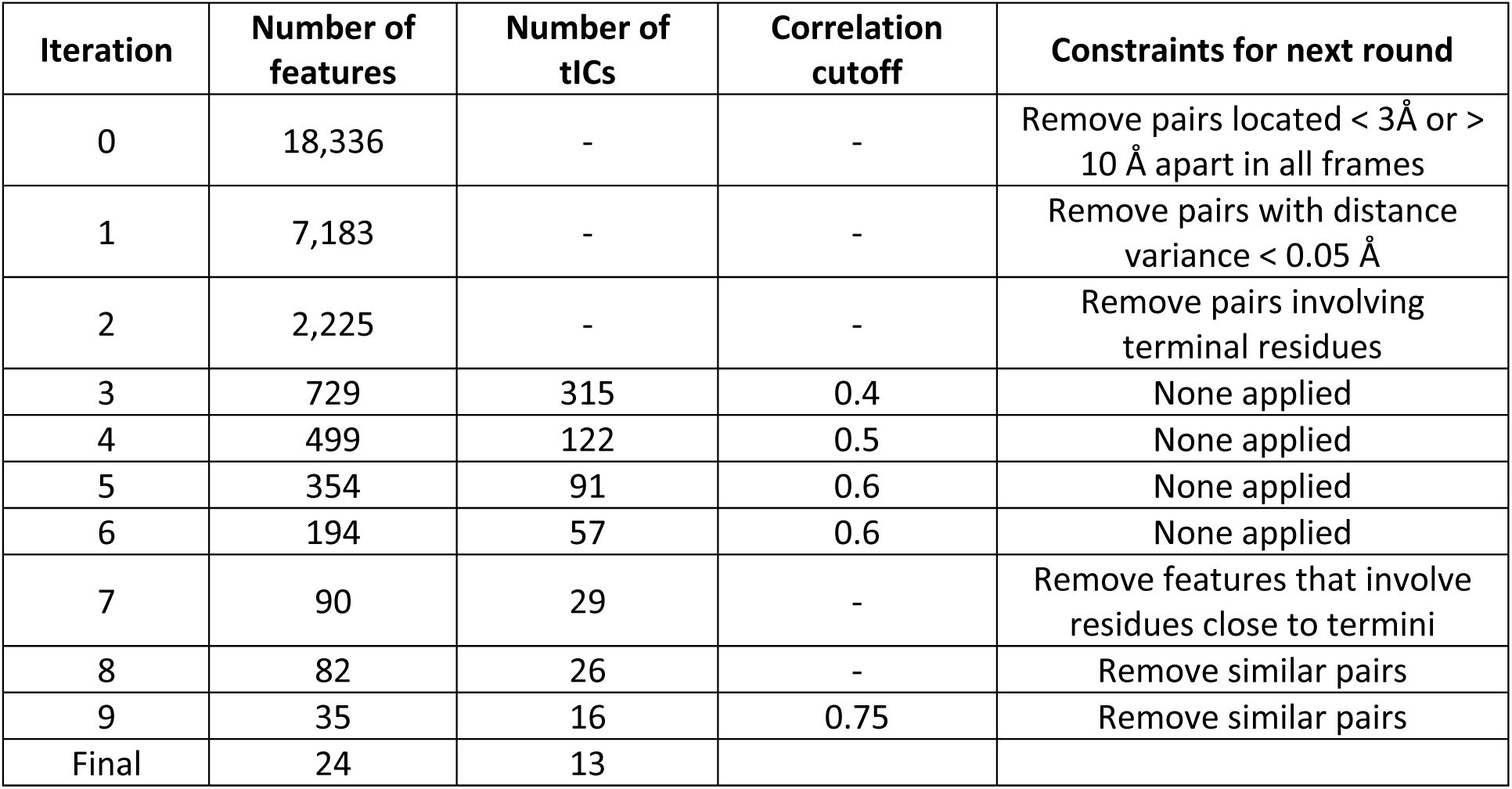
Stepwise tICA-based selection of features for model building.

**Supplementary Table 2.**
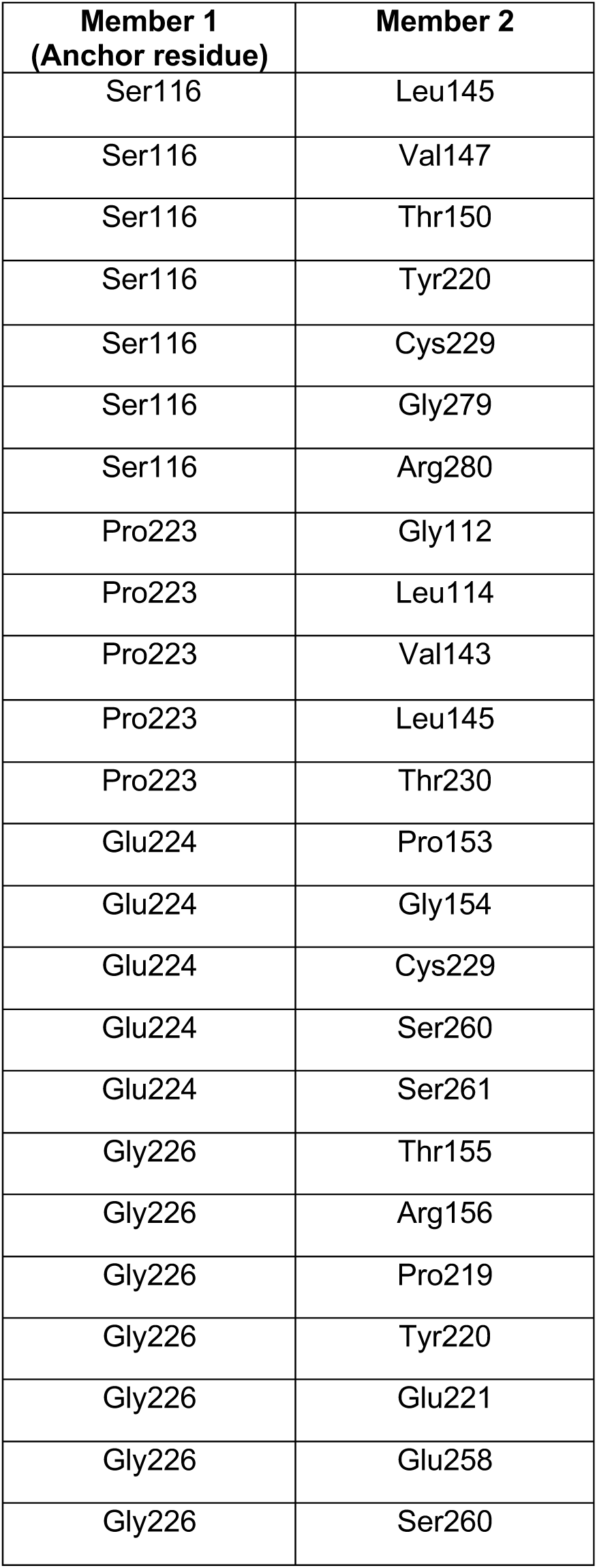
Pairs used for featurization of the simulations for model construction

**Supplementary Table 3.**
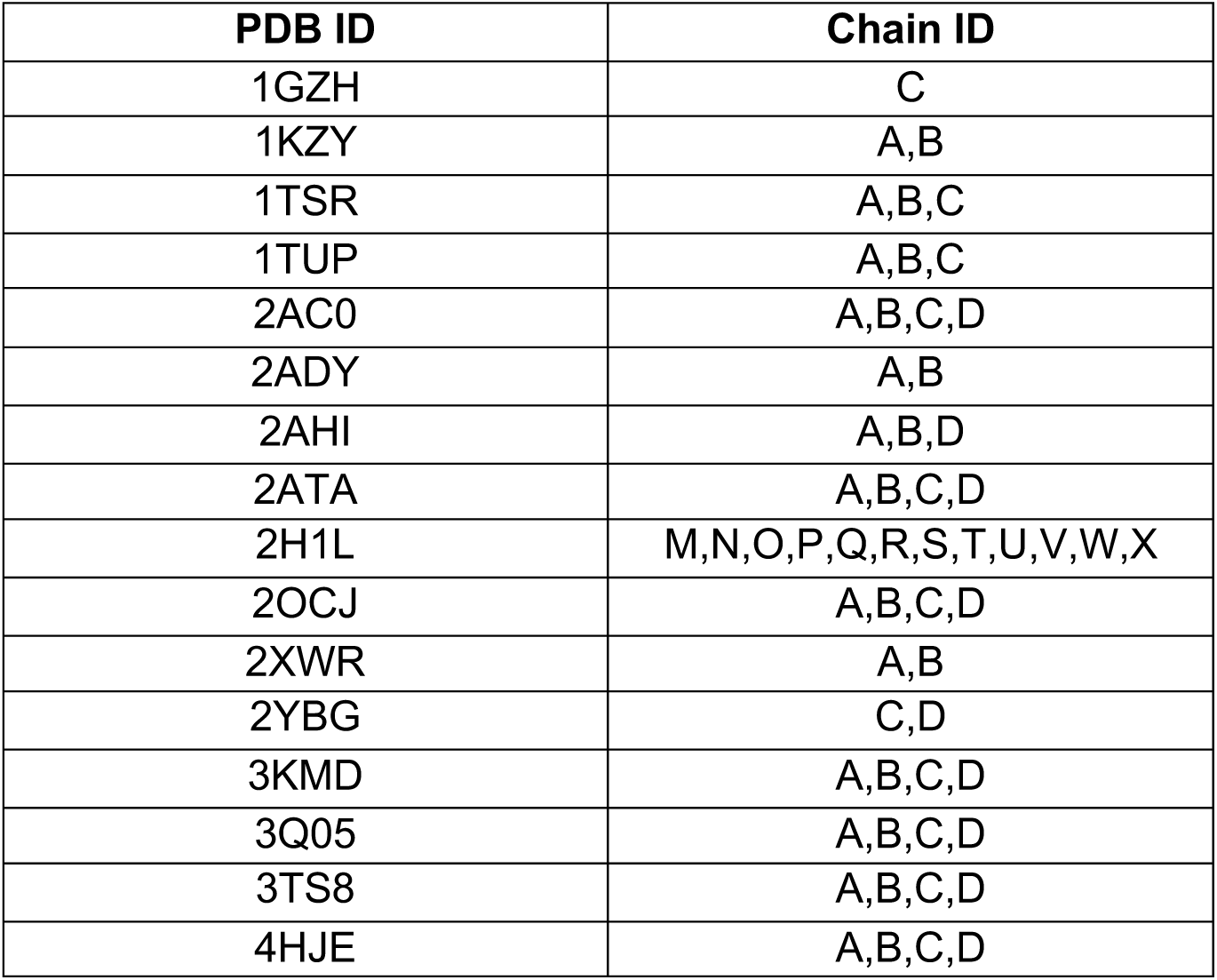
Wildtype X-ray and NMR structures used for comparison with simulations’ conformational landscape

**Supplementary Table 4.**
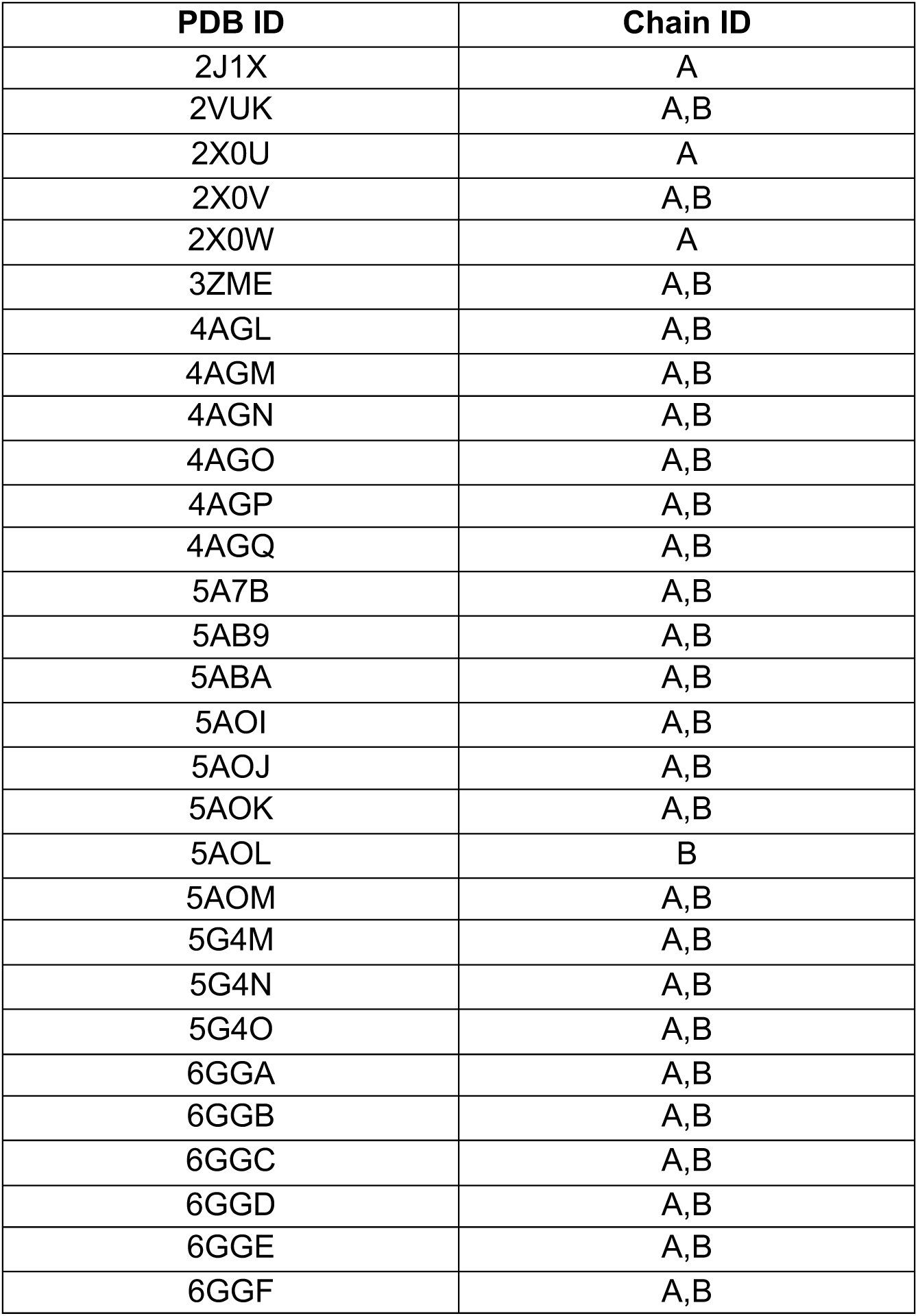
Y220C X-ray and NMR structures used for comparison with simulations’ conformational landscape

**Supplementary Table 5.**
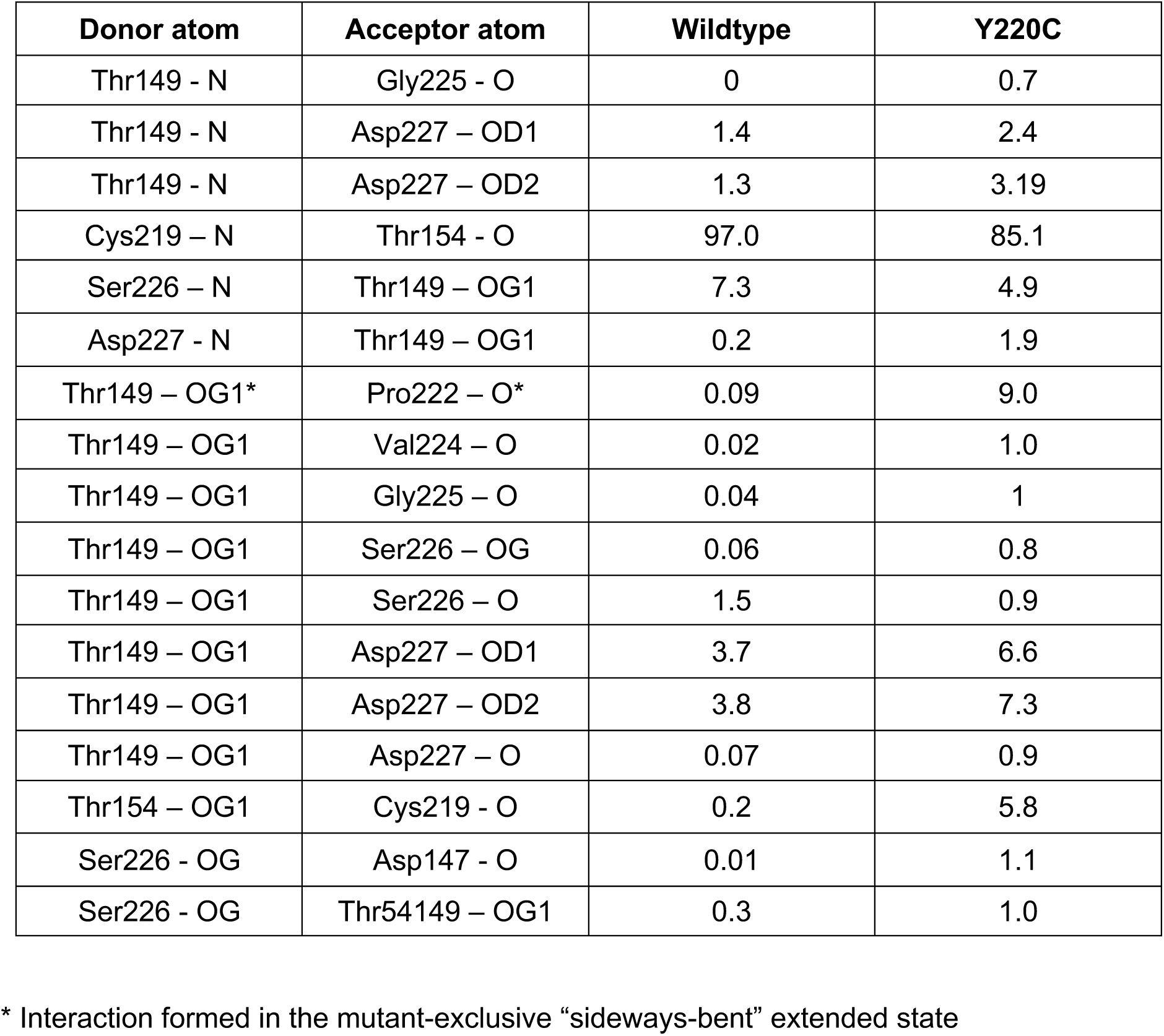
Persistence of L6-S3/S4 hydrogen bonds (in % of frames in the simulation)

**Supplementary Table 6.**
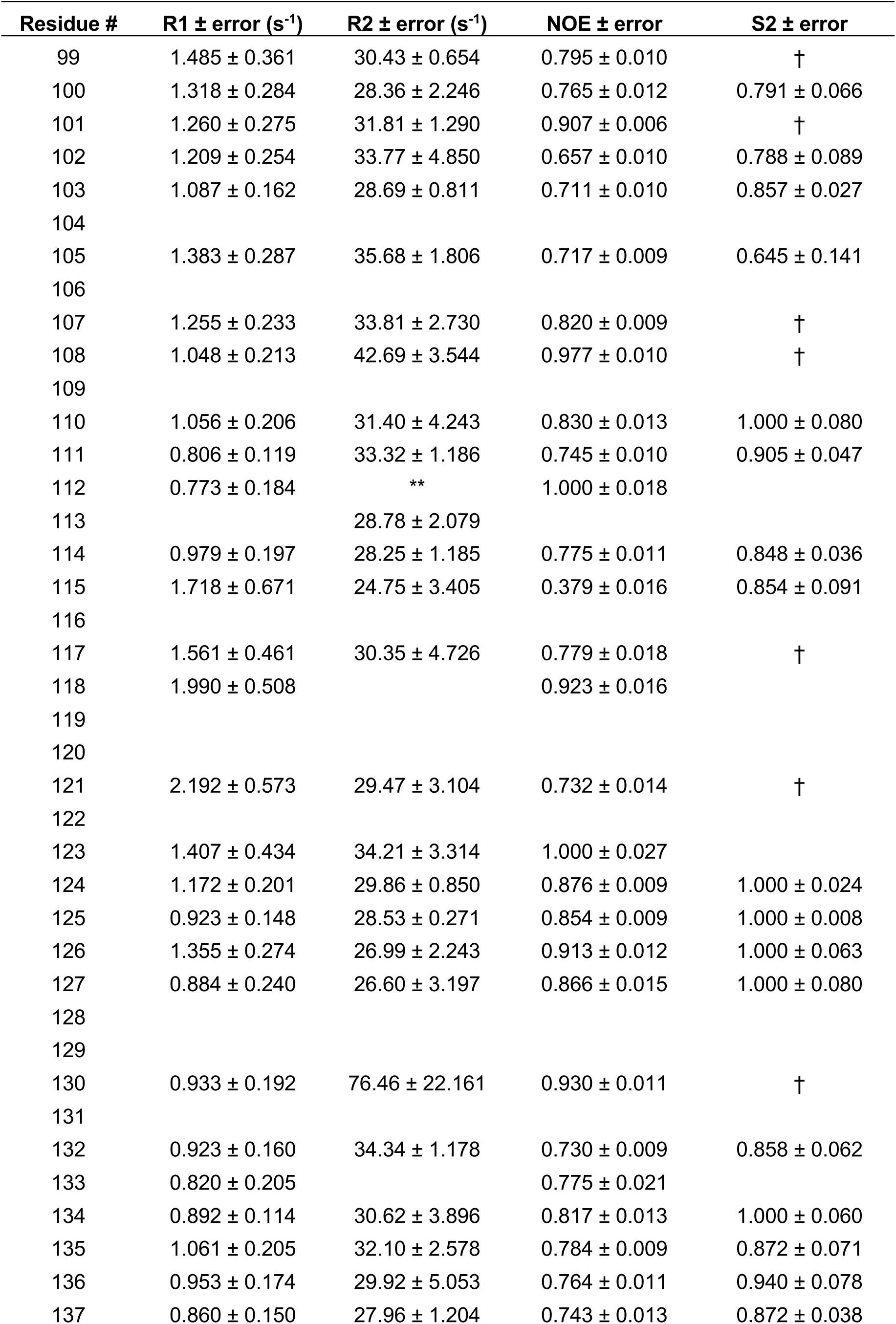
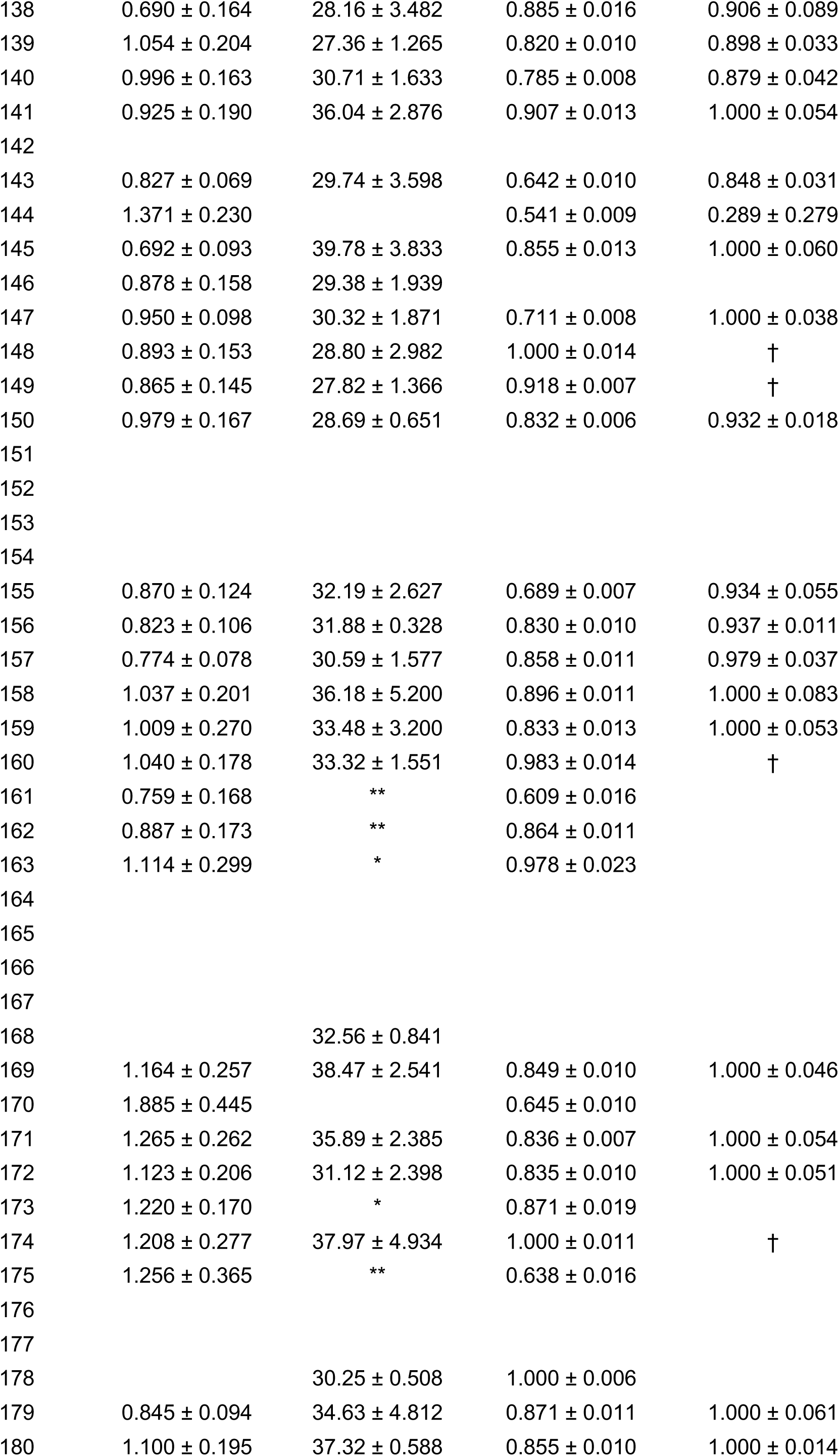
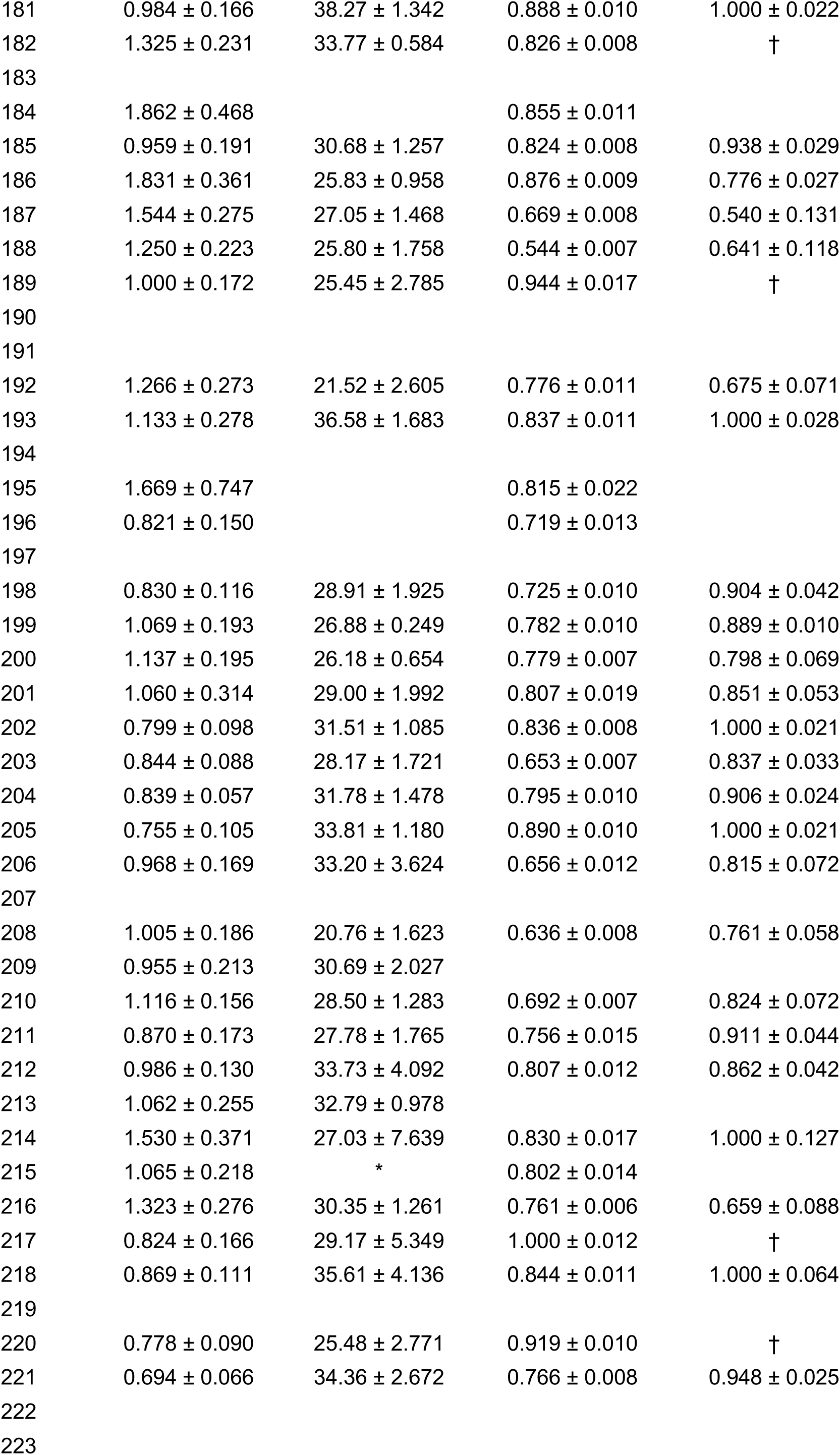
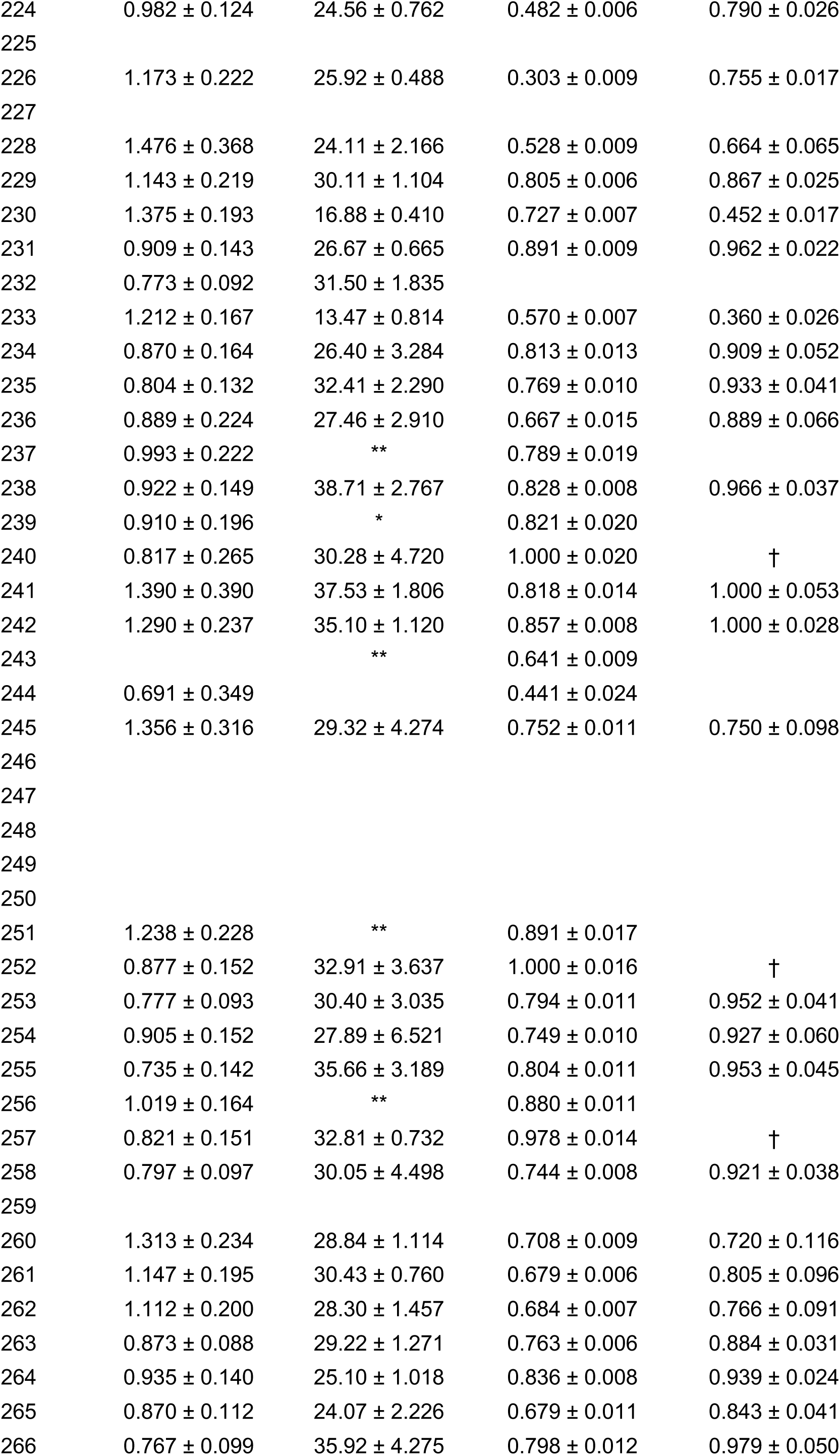
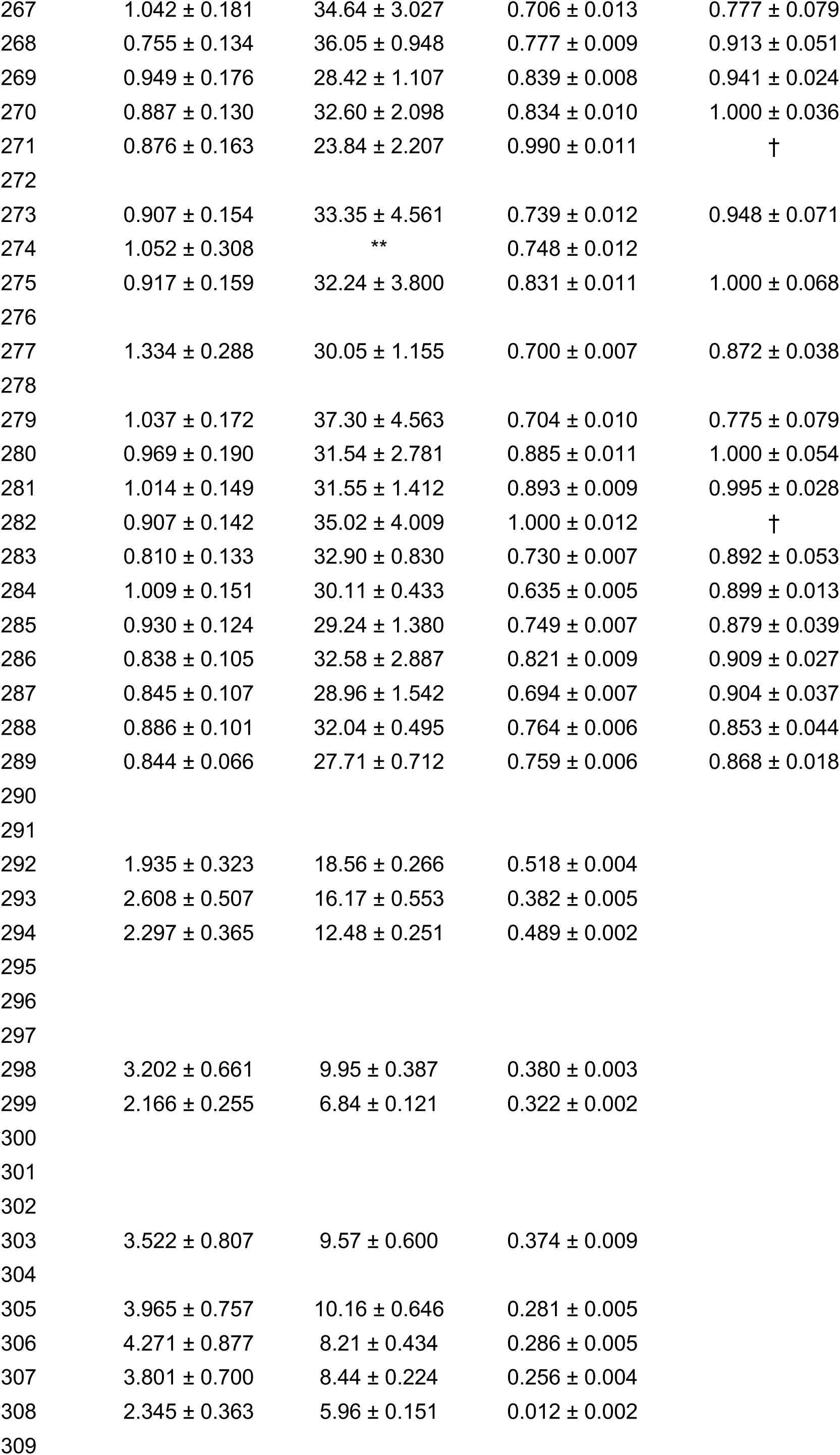
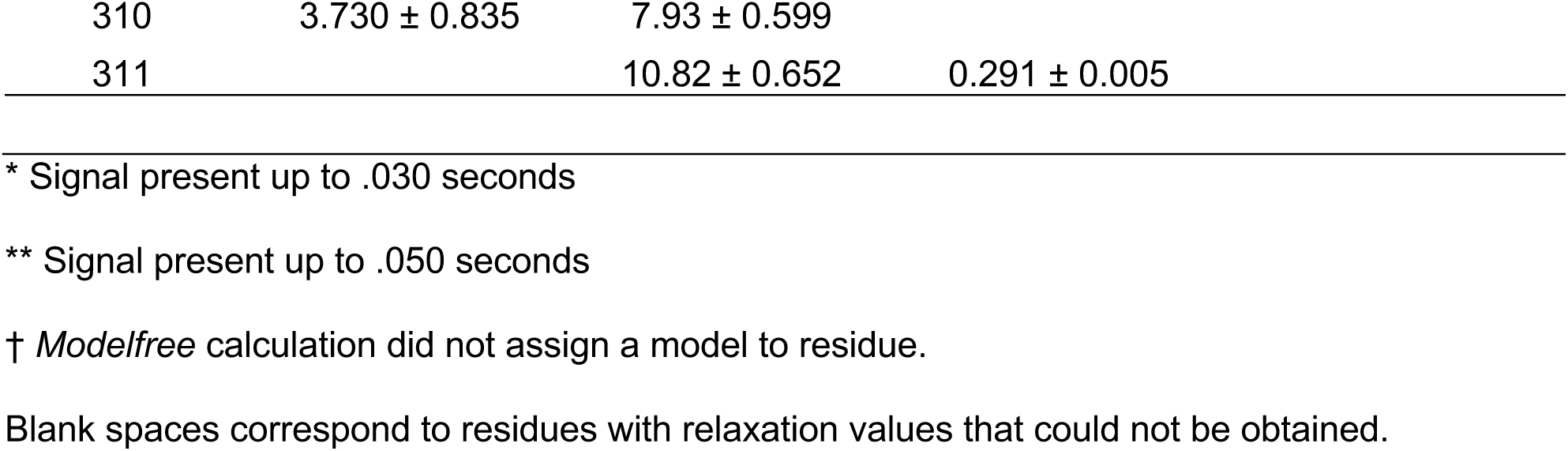
R1, R2, NOE, and S^2^ values for Wildtype p53 DBD at 800 MHz field strength.

| Residue # | R1 ± error (s<sup>-1</sup>) | R2 ± error (s<sup>-1</sup>) | NOE ± error | S2 ± error |
| --- | --- | --- | --- | --- |
| 99 | 1.485 ± 0.361 | 30.43 ± 0.654 | 0.795 ± 0.010 | † |
| 100 | 1.318 ± 0.284 | 28.36 ± 2.246 | 0.765 ± 0.012 | 0.791 ± 0.066 |
| 101 | 1.260 ± 0.275 | 31.81 ± 1.290 | 0.907 ± 0.006 | † |
| 102 | 1.209 ± 0.254 | 33.77 ± 4.850 | 0.657 ± 0.010 | 0.788 ± 0.089 |
| 103 | 1.087 ± 0.162 | 28.69 ± 0.811 | 0.711 ± 0.010 | 0.857 ± 0.027 |
| 104 |  |  |  |  |
| 105 | 1.383 ± 0.287 | 35.68 ± 1.806 | 0.717 ± 0.009 | 0.645 ± 0.141 |
| 106 |  |  |  |  |
| 107 | 1.255 ± 0.233 | 33.81 ± 2.730 | 0.820 ± 0.009 | † |
| 108 | 1.048 ± 0.213 | 42.69 ± 3.544 | 0.977 ± 0.010 | † |
| 109 |  |  |  |  |
| 110 | 1.056 ± 0.206 | 31.40 ± 4.243 | 0.830 ± 0.013 | 1.000 ± 0.080 |
| 111 | 0.806 ± 0.119 | 33.32 ± 1.186 | 0.745 ± 0.010 | 0.905 ± 0.047 |
| 112 | 0.773 ± 0.184 | ** | 1.000 ± 0.018 |  |
| 113 |  | 28.78 ± 2.079 |  |  |
| 114 | 0.979 ± 0.197 | 28.25 ± 1.185 | 0.775 ± 0.011 | 0.848 ± 0.036 |
| 115 | 1.718 ± 0.671 | 24.75 ± 3.405 | 0.379 ± 0.016 | 0.854 ± 0.091 |
| 116 |  |  |  |  |
| 117 | 1.561 ± 0.461 | 30.35 ± 4.726 | 0.779 ± 0.018 | † |
| 118 | 1.990 ± 0.508 |  | 0.923 ± 0.016 |  |
| 119 |  |  |  |  |
| 120 |  |  |  |  |
| 121 | 2.192 ± 0.573 | 29.47 ± 3.104 | 0.732 ± 0.014 | † |
| 122 |  |  |  |  |
| 123 | 1.407 ± 0.434 | 34.21 ± 3.314 | 1.000 ± 0.027 |  |
| 124 | 1.172 ± 0.201 | 29.86 ± 0.850 | 0.876 ± 0.009 | 1.000 ± 0.024 |
| 125 | 0.923 ± 0.148 | 28.53 ± 0.271 | 0.854 ± 0.009 | 1.000 ± 0.008 |
| 126 | 1.355 ± 0.274 | 26.99 ± 2.243 | 0.913 ± 0.012 | 1.000 ± 0.063 |
| 127 | 0.884 ± 0.240 | 26.60 ± 3.197 | 0.866 ± 0.015 | 1.000 ± 0.080 |
| 128 |  |  |  |  |
| 129 |  |  |  |  |
| 130 | 0.933 ± 0.192 | 76.46 ± 22.161 | 0.930 ± 0.011 | † |
| 131 |  |  |  |  |
| 132 | 0.923 ± 0.160 | 34.34 ± 1.178 | 0.730 ± 0.009 | 0.858 ± 0.062 |
| 133 | 0.820 ± 0.205 |  | 0.775 ± 0.021 |  |
| 134 | 0.892 ± 0.114 | 30.62 ± 3.896 | 0.817 ± 0.013 | 1.000 ± 0.060 |
| 135 | 1.061 ± 0.205 | 32.10 ± 2.578 | 0.784 ± 0.009 | 0.872 ± 0.071 |
| 136 | 0.953 ± 0.174 | 29.92 ± 5.053 | 0.764 ± 0.011 | 0.940 ± 0.078 |
| 137 | 0.860 ± 0.150 | 27.96 ± 1.204 | 0.743 ± 0.013 | 0.872 ± 0.038 |
| 138 | $0.690 \pm 0.164$ | $28.16 \pm 3.482$ | $0.885 \pm 0.016$ | $0.906 \pm 0.089$ |
| 139 | $1.054 \pm 0.204$ | $27.36 \pm 1.265$ | $0.820 \pm 0.010$ | $0.898 \pm 0.033$ |
| 140 | $0.996 \pm 0.163$ | $30.71 \pm 1.633$ | $0.785 \pm 0.008$ | $0.879 \pm 0.042$ |
| 141 | $0.925 \pm 0.190$ | $36.04 \pm 2.876$ | $0.907 \pm 0.013$ | $1.000 \pm 0.054$ |
| 142 |  |  |  |  |
| 143 | $0.827 \pm 0.069$ | $29.74 \pm 3.598$ | $0.642 \pm 0.010$ | $0.848 \pm 0.031$ |
| 144 | $1.371 \pm 0.230$ | | $0.541 \pm 0.009$ | $0.289 \pm 0.279$ |
| 145 | $0.692 \pm 0.093$ | $39.78 \pm 3.833$ | $0.855 \pm 0.013$ | $1.000 \pm 0.060$ |
| 146 | $0.878 \pm 0.158$ | $29.38 \pm 1.939$ | | |
| 147 | $0.950 \pm 0.098$ | $30.32 \pm 1.871$ | $0.711 \pm 0.008$ | $1.000 \pm 0.038$ |
| 148 | $0.893 \pm 0.153$ | $28.80 \pm 2.982$ | $1.000 \pm 0.014$ | † |
| 149 | $0.865 \pm 0.145$ | $27.82 \pm 1.366$ | $0.918 \pm 0.007$ | † |
| 150 | $0.979 \pm 0.167$ | $28.69 \pm 0.651$ | $0.832 \pm 0.006$ | $0.932 \pm 0.018$ |
| 151 |  |  |  |  |
| 152 |  |  |  |  |
| 153 |  |  |  |  |
| 154 |  |  |  |  |
| 155 | $0.870 \pm 0.124$ | $32.19 \pm 2.627$ | $0.689 \pm 0.007$ | $0.934 \pm 0.055$ |
| 156 | $0.823 \pm 0.106$ | $31.88 \pm 0.328$ | $0.830 \pm 0.010$ | $0.937 \pm 0.011$ |
| 157 | $0.774 \pm 0.078$ | $30.59 \pm 1.577$ | $0.858 \pm 0.011$ | $0.979 \pm 0.037$ |
| 158 | $1.037 \pm 0.201$ | $36.18 \pm 5.200$ | $0.896 \pm 0.011$ | $1.000 \pm 0.083$ |
| 159 | $1.009 \pm 0.270$ | $33.48 \pm 3.200$ | $0.833 \pm 0.013$ | $1.000 \pm 0.053$ |
| 160 | $1.040 \pm 0.178$ | $33.32 \pm 1.551$ | $0.983 \pm 0.014$ | † |
| 161 | $0.759 \pm 0.168$ | ** | $0.609 \pm 0.016$ | |
| 162 | $0.887 \pm 0.173$ | ** | $0.864 \pm 0.011$ | |
| 163 | $1.114 \pm 0.299$ | * | $0.978 \pm 0.023$ | |
| 164 |  |  |  |  |
| 165 |  |  |  |  |
| 166 |  |  |  |  |
| 167 |  |  |  |  |
| 168 | | $32.56 \pm 0.841$ | | |
| 169 | $1.164 \pm 0.257$ | $38.47 \pm 2.541$ | $0.849 \pm 0.010$ | $1.000 \pm 0.046$ |
| 170 | $1.885 \pm 0.445$ | | $0.645 \pm 0.010$ | |
| 171 | $1.265 \pm 0.262$ | $35.89 \pm 2.385$ | $0.836 \pm 0.007$ | $1.000 \pm 0.054$ |
| 172 | $1.123 \pm 0.206$ | $31.12 \pm 2.398$ | $0.835 \pm 0.010$ | $1.000 \pm 0.051$ |
| 173 | $1.220 \pm 0.170$ | * | $0.871 \pm 0.019$ | |
| 174 | $1.208 \pm 0.277$ | $37.97 \pm 4.934$ | $1.000 \pm 0.011$ | † |
| 175 | $1.256 \pm 0.365$ | ** | $0.638 \pm 0.016$ | |
| 176 |  |  |  |  |
| 177 |  |  |  |  |
| 178 | | $30.25 \pm 0.508$ | $1.000 \pm 0.006$ | |
| 179 | $0.845 \pm 0.094$ | $34.63 \pm 4.812$ | $0.871 \pm 0.011$ | $1.000 \pm 0.061$ |
| 180 | $1.100 \pm 0.195$ | $37.32 \pm 0.588$ | $0.855 \pm 0.010$ | $1.000 \pm 0.014$ |
| 181 | $0.984 \pm 0.166$ | $38.27 \pm 1.342$ | $0.888 \pm 0.010$ | $1.000 \pm 0.022$ |
| 182 | $1.325 \pm 0.231$ | $33.77 \pm 0.584$ | $0.826 \pm 0.008$ | † |
| 183 |  |  |  |  |
| 184 | $1.862 \pm 0.468$ | | $0.855 \pm 0.011$ | |
| 185 | $0.959 \pm 0.191$ | $30.68 \pm 1.257$ | $0.824 \pm 0.008$ | $0.938 \pm 0.029$ |
| 186 | $1.831 \pm 0.361$ | $25.83 \pm 0.958$ | $0.876 \pm 0.009$ | $0.776 \pm 0.027$ |
| 187 | $1.544 \pm 0.275$ | $27.05 \pm 1.468$ | $0.669 \pm 0.008$ | $0.540 \pm 0.131$ |
| 188 | $1.250 \pm 0.223$ | $25.80 \pm 1.758$ | $0.544 \pm 0.007$ | $0.641 \pm 0.118$ |
| 189 | $1.000 \pm 0.172$ | $25.45 \pm 2.785$ | $0.944 \pm 0.017$ | † |
| 190 |  |  |  |  |
| 191 |  |  |  |  |
| 192 | $1.266 \pm 0.273$ | $21.52 \pm 2.605$ | $0.776 \pm 0.011$ | $0.675 \pm 0.071$ |
| 193 | $1.133 \pm 0.278$ | $36.58 \pm 1.683$ | $0.837 \pm 0.011$ | $1.000 \pm 0.028$ |
| 194 |  |  |  |  |
| 195 | $1.669 \pm 0.747$ | | $0.815 \pm 0.022$ | |
| 196 | $0.821 \pm 0.150$ | | $0.719 \pm 0.013$ | |
| 197 |  |  |  |  |
| 198 | $0.830 \pm 0.116$ | $28.91 \pm 1.925$ | $0.725 \pm 0.010$ | $0.904 \pm 0.042$ |
| 199 | $1.069 \pm 0.193$ | $26.88 \pm 0.249$ | $0.782 \pm 0.010$ | $0.889 \pm 0.010$ |
| 200 | $1.137 \pm 0.195$ | $26.18 \pm 0.654$ | $0.779 \pm 0.007$ | $0.798 \pm 0.069$ |
| 201 | $1.060 \pm 0.314$ | $29.00 \pm 1.992$ | $0.807 \pm 0.019$ | $0.851 \pm 0.053$ |
| 202 | $0.799 \pm 0.098$ | $31.51 \pm 1.085$ | $0.836 \pm 0.008$ | $1.000 \pm 0.021$ |
| 203 | $0.844 \pm 0.088$ | $28.17 \pm 1.721$ | $0.653 \pm 0.007$ | $0.837 \pm 0.033$ |
| 204 | $0.839 \pm 0.057$ | $31.78 \pm 1.478$ | $0.795 \pm 0.010$ | $0.906 \pm 0.024$ |
| 205 | $0.755 \pm 0.105$ | $33.81 \pm 1.180$ | $0.890 \pm 0.010$ | $1.000 \pm 0.021$ |
| 206 | $0.968 \pm 0.169$ | $33.20 \pm 3.624$ | $0.656 \pm 0.012$ | $0.815 \pm 0.072$ |
| 207 |  |  |  |  |
| 208 | $1.005 \pm 0.186$ | $20.76 \pm 1.623$ | $0.636 \pm 0.008$ | $0.761 \pm 0.058$ |
| 209 | $0.955 \pm 0.213$ | $30.69 \pm 2.027$ | | |
| 210 | $1.116 \pm 0.156$ | $28.50 \pm 1.283$ | $0.692 \pm 0.007$ | $0.824 \pm 0.072$ |
| 211 | $0.870 \pm 0.173$ | $27.78 \pm 1.765$ | $0.756 \pm 0.015$ | $0.911 \pm 0.044$ |
| 212 | $0.986 \pm 0.130$ | $33.73 \pm 4.092$ | $0.807 \pm 0.012$ | $0.862 \pm 0.042$ |
| 213 | $1.062 \pm 0.255$ | $32.79 \pm 0.978$ | | |
| 214 | $1.530 \pm 0.371$ | $27.03 \pm 7.639$ | $0.830 \pm 0.017$ | $1.000 \pm 0.127$ |
| 215 | $1.065 \pm 0.218$ | * | $0.802 \pm 0.014$ | |
| 216 | $1.323 \pm 0.276$ | $30.35 \pm 1.261$ | $0.761 \pm 0.006$ | $0.659 \pm 0.088$ |
| 217 | $0.824 \pm 0.166$ | $29.17 \pm 5.349$ | $1.000 \pm 0.012$ | † |
| 218 | $0.869 \pm 0.111$ | $35.61 \pm 4.136$ | $0.844 \pm 0.011$ | $1.000 \pm 0.064$ |
| 219 |  |  |  |  |
| 220 | $0.778 \pm 0.090$ | $25.48 \pm 2.771$ | $0.919 \pm 0.010$ | † |
| 221 | $0.694 \pm 0.066$ | $34.36 \pm 2.672$ | $0.766 \pm 0.008$ | $0.948 \pm 0.025$ |
| 222 |  |  |  |  |
| 223 |  |  |  |  |
| 224 | 0.982 ± 0.124 | 24.56 ± 0.762 | 0.482 ± 0.006 | 0.790 ± 0.026 |
| 225 |  |  |  |  |
| 226 | 1.173 ± 0.222 | 25.92 ± 0.488 | 0.303 ± 0.009 | 0.755 ± 0.017 |
| 227 |  |  |  |  |
| 228 | 1.476 ± 0.368 | 24.11 ± 2.166 | 0.528 ± 0.009 | 0.664 ± 0.065 |
| 229 | 1.143 ± 0.219 | 30.11 ± 1.104 | 0.805 ± 0.006 | 0.867 ± 0.025 |
| 230 | 1.375 ± 0.193 | 16.88 ± 0.410 | 0.727 ± 0.007 | 0.452 ± 0.017 |
| 231 | 0.909 ± 0.143 | 26.67 ± 0.665 | 0.891 ± 0.009 | 0.962 ± 0.022 |
| 232 | 0.773 ± 0.092 | 31.50 ± 1.835 |  |  |
| 233 | 1.212 ± 0.167 | 13.47 ± 0.814 | 0.570 ± 0.007 | 0.360 ± 0.026 |
| 234 | 0.870 ± 0.164 | 26.40 ± 3.284 | 0.813 ± 0.013 | 0.909 ± 0.052 |
| 235 | 0.804 ± 0.132 | 32.41 ± 2.290 | 0.769 ± 0.010 | 0.933 ± 0.041 |
| 236 | 0.889 ± 0.224 | 27.46 ± 2.910 | 0.667 ± 0.015 | 0.889 ± 0.066 |
| 237 | 0.993 ± 0.222 | ** | 0.789 ± 0.019 |  |
| 238 | 0.922 ± 0.149 | 38.71 ± 2.767 | 0.828 ± 0.008 | 0.966 ± 0.037 |
| 239 | 0.910 ± 0.196 | * | 0.821 ± 0.020 |  |
| 240 | 0.817 ± 0.265 | 30.28 ± 4.720 | 1.000 ± 0.020 | † |
| 241 | 1.390 ± 0.390 | 37.53 ± 1.806 | 0.818 ± 0.014 | 1.000 ± 0.053 |
| 242 | 1.290 ± 0.237 | 35.10 ± 1.120 | 0.857 ± 0.008 | 1.000 ± 0.028 |
| 243 |  | ** | 0.641 ± 0.009 |  |
| 244 | 0.691 ± 0.349 |  | 0.441 ± 0.024 |  |
| 245 | 1.356 ± 0.316 | 29.32 ± 4.274 | 0.752 ± 0.011 | 0.750 ± 0.098 |
| 246 |  |  |  |  |
| 247 |  |  |  |  |
| 248 |  |  |  |  |
| 249 |  |  |  |  |
| 250 |  |  |  |  |
| 251 | 1.238 ± 0.228 | ** | 0.891 ± 0.017 |  |
| 252 | 0.877 ± 0.152 | 32.91 ± 3.637 | 1.000 ± 0.016 | † |
| 253 | 0.777 ± 0.093 | 30.40 ± 3.035 | 0.794 ± 0.011 | 0.952 ± 0.041 |
| 254 | 0.905 ± 0.152 | 27.89 ± 6.521 | 0.749 ± 0.010 | 0.927 ± 0.060 |
| 255 | 0.735 ± 0.142 | 35.66 ± 3.189 | 0.804 ± 0.011 | 0.953 ± 0.045 |
| 256 | 1.019 ± 0.164 | ** | 0.880 ± 0.011 |  |
| 257 | 0.821 ± 0.151 | 32.81 ± 0.732 | 0.978 ± 0.014 | † |
| 258 | 0.797 ± 0.097 | 30.05 ± 4.498 | 0.744 ± 0.008 | 0.921 ± 0.038 |
| 259 |  |  |  |  |
| 260 | 1.313 ± 0.234 | 28.84 ± 1.114 | 0.708 ± 0.009 | 0.720 ± 0.116 |
| 261 | 1.147 ± 0.195 | 30.43 ± 0.760 | 0.679 ± 0.006 | 0.805 ± 0.096 |
| 262 | 1.112 ± 0.200 | 28.30 ± 1.457 | 0.684 ± 0.007 | 0.766 ± 0.091 |
| 263 | 0.873 ± 0.088 | 29.22 ± 1.271 | 0.763 ± 0.006 | 0.884 ± 0.031 |
| 264 | 0.935 ± 0.140 | 25.10 ± 1.018 | 0.836 ± 0.008 | 0.939 ± 0.024 |
| 265 | 0.870 ± 0.112 | 24.07 ± 2.226 | 0.679 ± 0.011 | 0.843 ± 0.041 |
| 266 | 0.767 ± 0.099 | 35.92 ± 4.275 | 0.798 ± 0.012 | 0.979 ± 0.050 |
| 267 | $1.042 \pm 0.181$ | $34.64 \pm 3.027$ | $0.706 \pm 0.013$ | $0.777 \pm 0.079$ |
| 268 | $0.755 \pm 0.134$ | $36.05 \pm 0.948$ | $0.777 \pm 0.009$ | $0.913 \pm 0.051$ |
| 269 | $0.949 \pm 0.176$ | $28.42 \pm 1.107$ | $0.839 \pm 0.008$ | $0.941 \pm 0.024$ |
| 270 | $0.887 \pm 0.130$ | $32.60 \pm 2.098$ | $0.834 \pm 0.010$ | $1.000 \pm 0.036$ |
| 271 | $0.876 \pm 0.163$ | $23.84 \pm 2.207$ | $0.990 \pm 0.011$ | † |
| 272 |  |  |  |  |
| 273 | $0.907 \pm 0.154$ | $33.35 \pm 4.561$ | $0.739 \pm 0.012$ | $0.948 \pm 0.071$ |
| 274 | $1.052 \pm 0.308$ | ** | $0.748 \pm 0.012$ | |
| 275 | $0.917 \pm 0.159$ | $32.24 \pm 3.800$ | $0.831 \pm 0.011$ | $1.000 \pm 0.068$ |
| 276 |  |  |  |  |
| 277 | $1.334 \pm 0.288$ | $30.05 \pm 1.155$ | $0.700 \pm 0.007$ | $0.872 \pm 0.038$ |
| 278 |  |  |  |  |
| 279 | $1.037 \pm 0.172$ | $37.30 \pm 4.563$ | $0.704 \pm 0.010$ | $0.775 \pm 0.079$ |
| 280 | $0.969 \pm 0.190$ | $31.54 \pm 2.781$ | $0.885 \pm 0.011$ | $1.000 \pm 0.054$ |
| 281 | $1.014 \pm 0.149$ | $31.55 \pm 1.412$ | $0.893 \pm 0.009$ | $0.995 \pm 0.028$ |
| 282 | $0.907 \pm 0.142$ | $35.02 \pm 4.009$ | $1.000 \pm 0.012$ | † |
| 283 | $0.810 \pm 0.133$ | $32.90 \pm 0.830$ | $0.730 \pm 0.007$ | $0.892 \pm 0.053$ |
| 284 | $1.009 \pm 0.151$ | $30.11 \pm 0.433$ | $0.635 \pm 0.005$ | $0.899 \pm 0.013$ |
| 285 | $0.930 \pm 0.124$ | $29.24 \pm 1.380$ | $0.749 \pm 0.007$ | $0.879 \pm 0.039$ |
| 286 | $0.838 \pm 0.105$ | $32.58 \pm 2.887$ | $0.821 \pm 0.009$ | $0.909 \pm 0.027$ |
| 287 | $0.845 \pm 0.107$ | $28.96 \pm 1.542$ | $0.694 \pm 0.007$ | $0.904 \pm 0.037$ |
| 288 | $0.886 \pm 0.101$ | $32.04 \pm 0.495$ | $0.764 \pm 0.006$ | $0.853 \pm 0.044$ |
| 289 | $0.844 \pm 0.066$ | $27.71 \pm 0.712$ | $0.759 \pm 0.006$ | $0.868 \pm 0.018$ |
| 290 |  |  |  |  |
| 291 |  |  |  |  |
| 292 | $1.935 \pm 0.323$ | $18.56 \pm 0.266$ | $0.518 \pm 0.004$ | |
| 293 | $2.608 \pm 0.507$ | $16.17 \pm 0.553$ | $0.382 \pm 0.005$ | |
| 294 | $2.297 \pm 0.365$ | $12.48 \pm 0.251$ | $0.489 \pm 0.002$ | |
| 295 |  |  |  |  |
| 296 |  |  |  |  |
| 297 |  |  |  |  |
| 298 | $3.202 \pm 0.661$ | $9.95 \pm 0.387$ | $0.380 \pm 0.003$ | |
| 299 | $2.166 \pm 0.255$ | $6.84 \pm 0.121$ | $0.322 \pm 0.002$ | |
| 300 |  |  |  |  |
| 301 |  |  |  |  |
| 302 |  |  |  |  |
| 303 | $3.522 \pm 0.807$ | $9.57 \pm 0.600$ | $0.374 \pm 0.009$ | |
| 304 |  |  |  |  |
| 305 | $3.965 \pm 0.757$ | $10.16 \pm 0.646$ | $0.281 \pm 0.005$ | |
| 306 | $4.271 \pm 0.877$ | $8.21 \pm 0.434$ | $0.286 \pm 0.005$ | |
| 307 | $3.801 \pm 0.700$ | $8.44 \pm 0.224$ | $0.256 \pm 0.004$ | |
| 308 | $2.345 \pm 0.363$ | $5.96 \pm 0.151$ | $0.012 \pm 0.002$ | |
| 309 |  |  |  |  |

|  |  |  |  |
| --- | --- | --- | --- |
| 310 | 3.730 ± 0.835 | 7.93 ± 0.599 |  |
| 311 |  | 10.82 ± 0.652 | 0.291 ± 0.005 |

\* Signal present up to .030 seconds
\*\* Signal present up to .050 seconds
† *Modelfree* calculation did not assign a model to residue.
Blank spaces correspond to residues with relaxation values that could not be obtained.

**Supplementary Figure 1.**
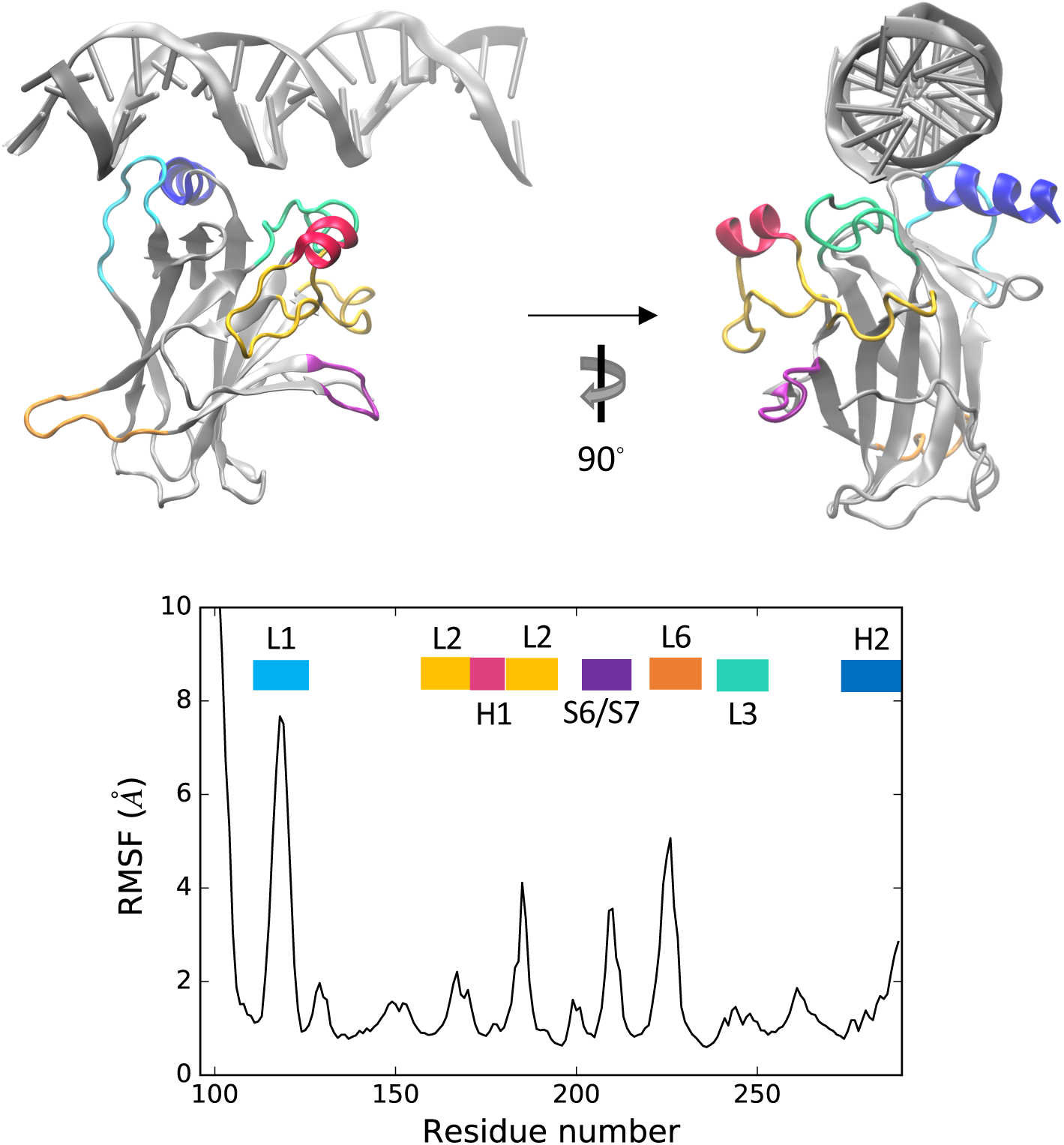
Alpha carbon RMSF. Functionally important structural motifs are highlighted in the DNA-bound structure (top panel).

**Supplementary Figure 2.**
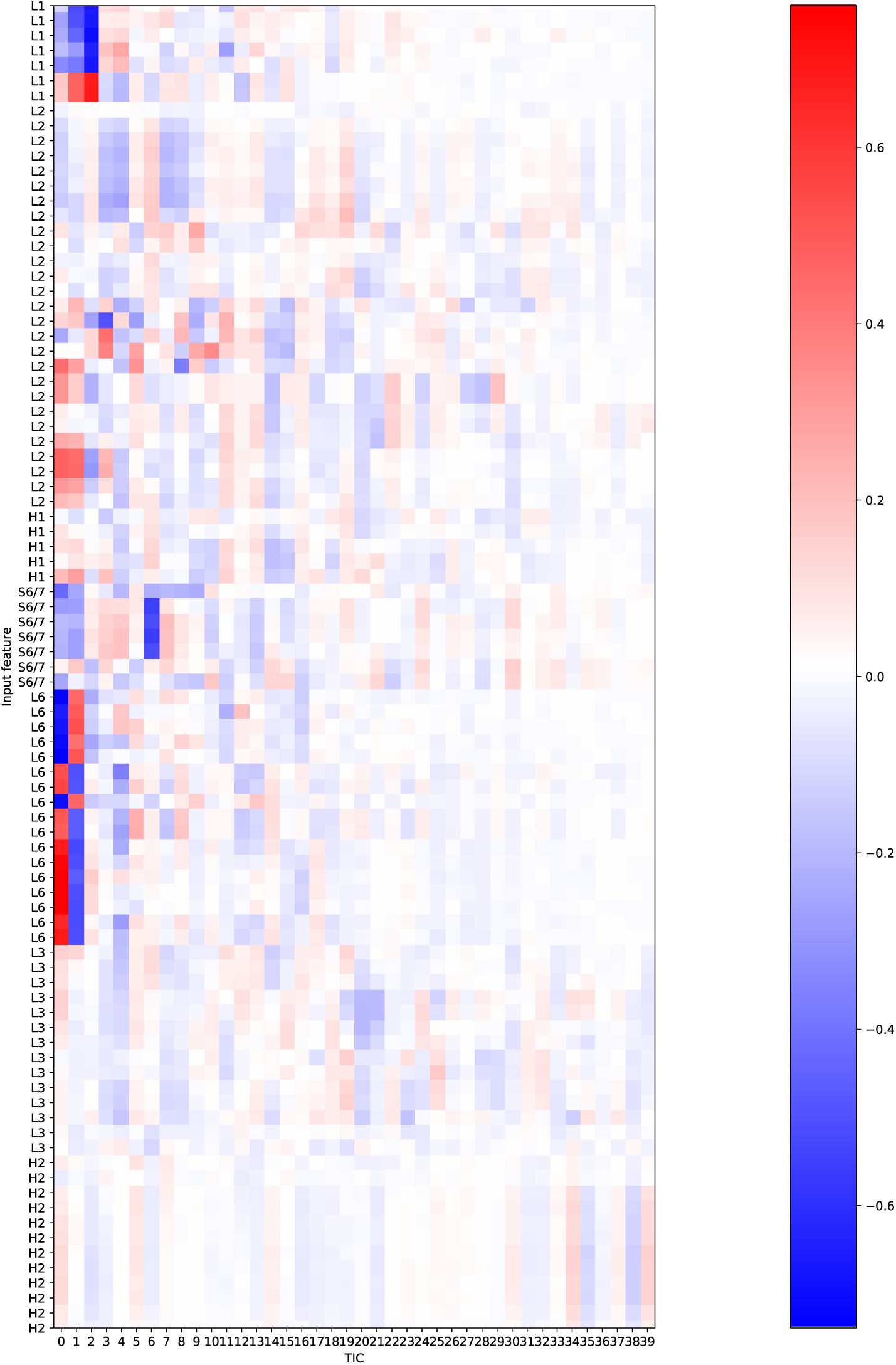
tICA correlation for features incorporating functionally-important motifs in the protein (H1, H2, L2, L3, S6/7) in addition to L1 and L6.

**Supplementary Figure 3.**
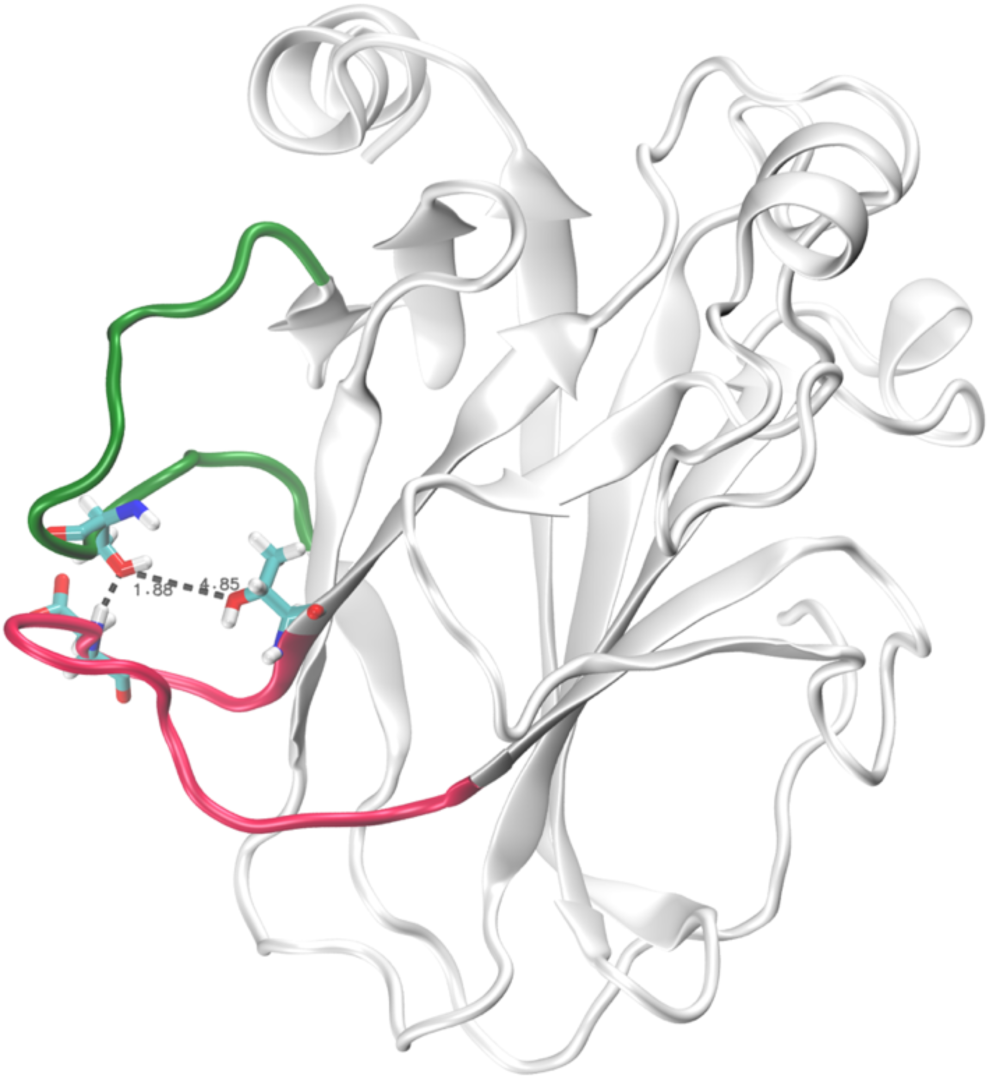
Example of frame exhibiting most stable intra-loop hydrogen bonds, involving Ser116 in L1 and Asp228 or Thr231 in L6.

**Supplementary Figure 4.**
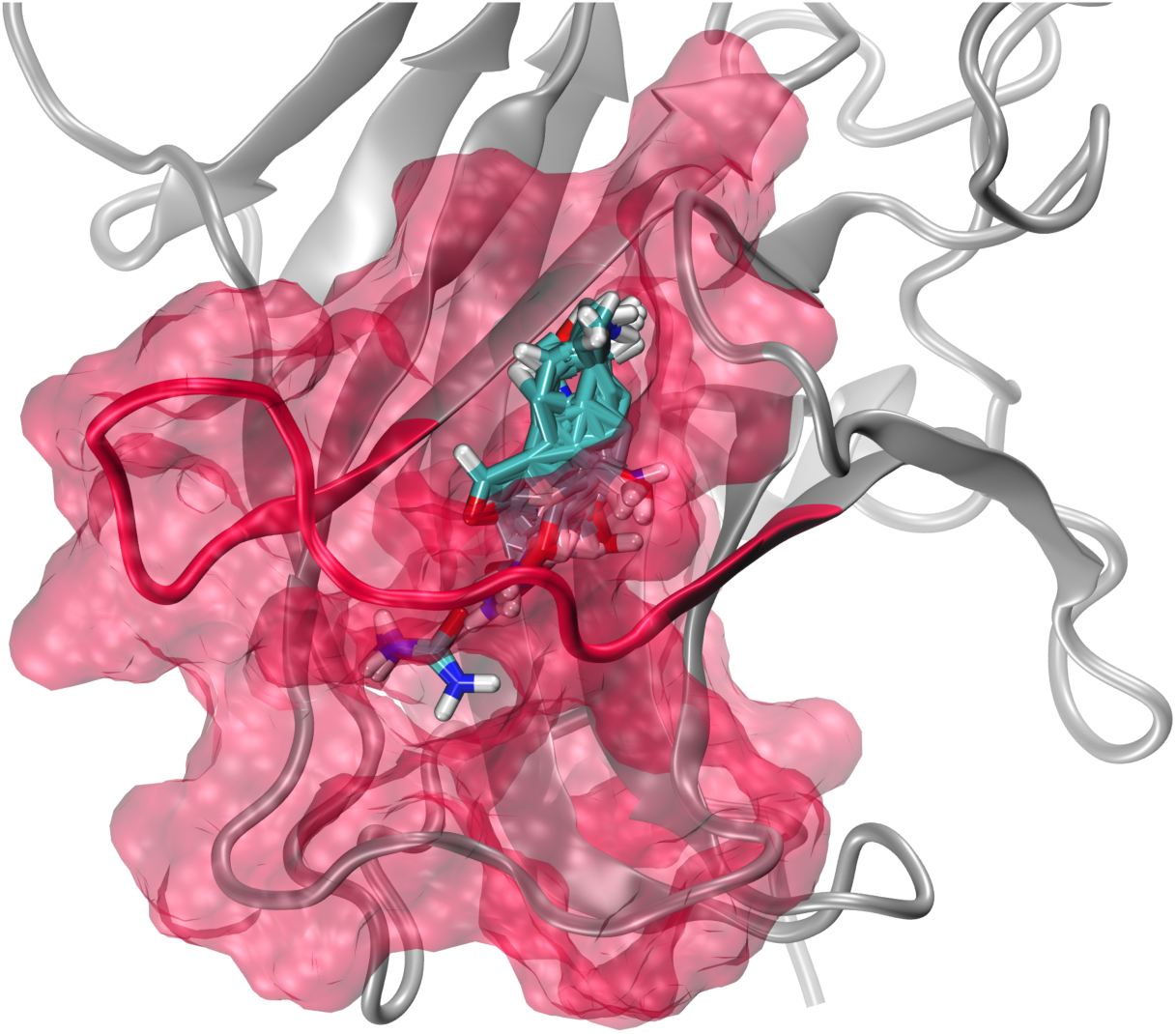
Representation of the cryptic channel spanning loop L6 in the recessed Y220C metastable state. FTMap^11^ probes indicating hotspots for drug binding are shown in licorice.

**Supplementary Figure 5.**
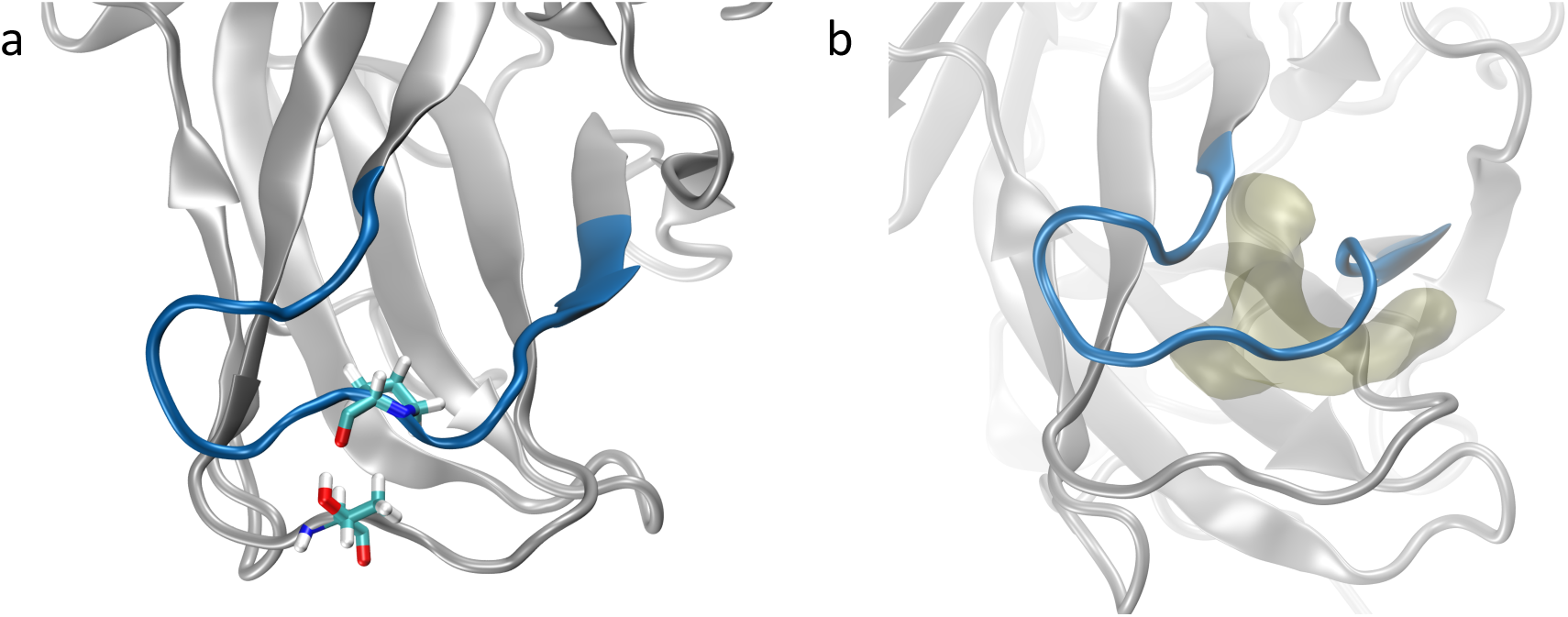
(a) Representation of the Thr149-Pro222 interaction thought to stabilize the bent L6 conformation observed in the mutant-exclusive states. (b) Surface representation of the L6 pocket in the mutant-exclusive states.

**Supplementary Figure 6.**
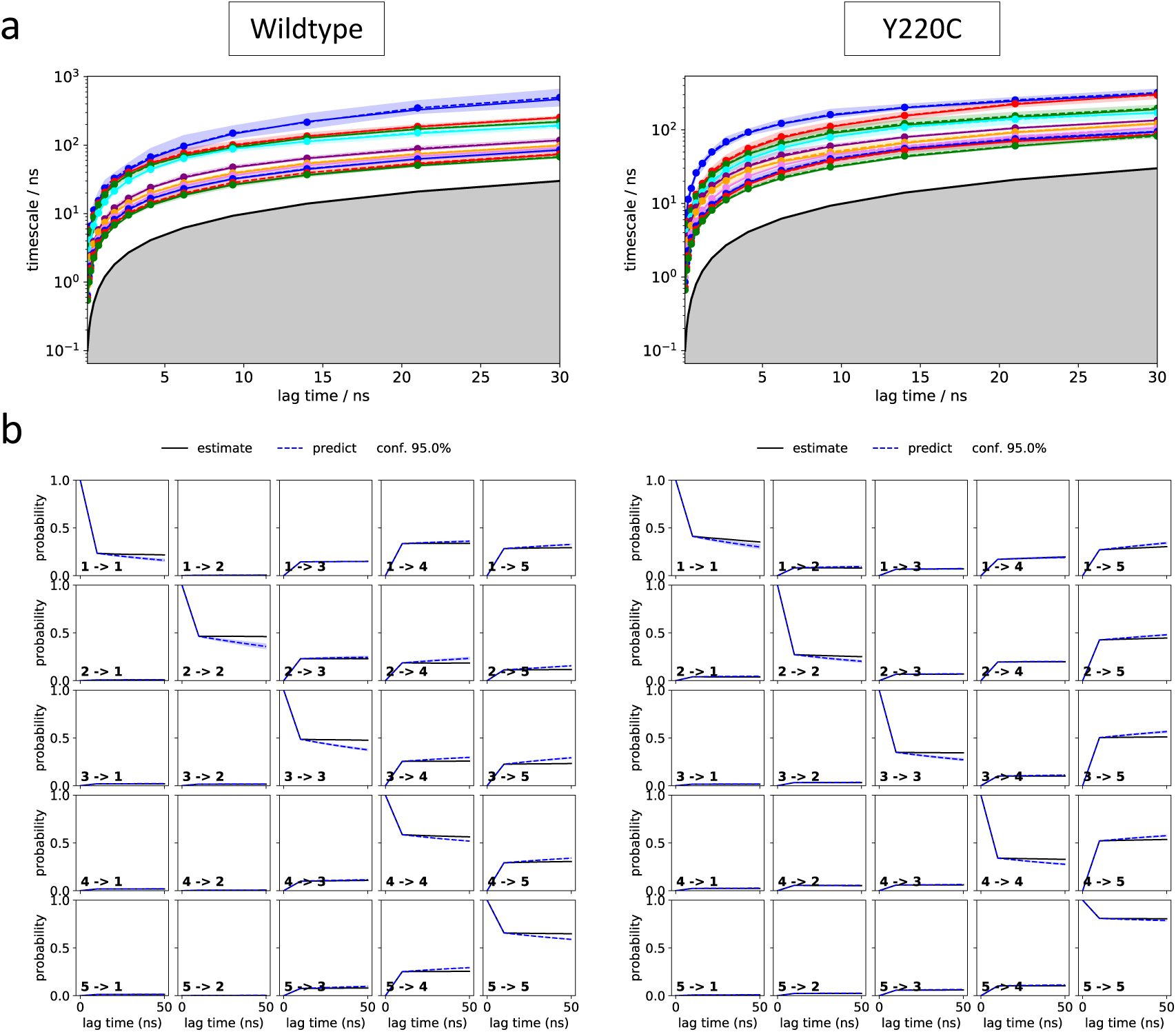
L1 MSM model validation analysis: (a) Implied timescale plots and (b) Chapman-Kolmogorov tests.

**Supplementary Figure 7.**
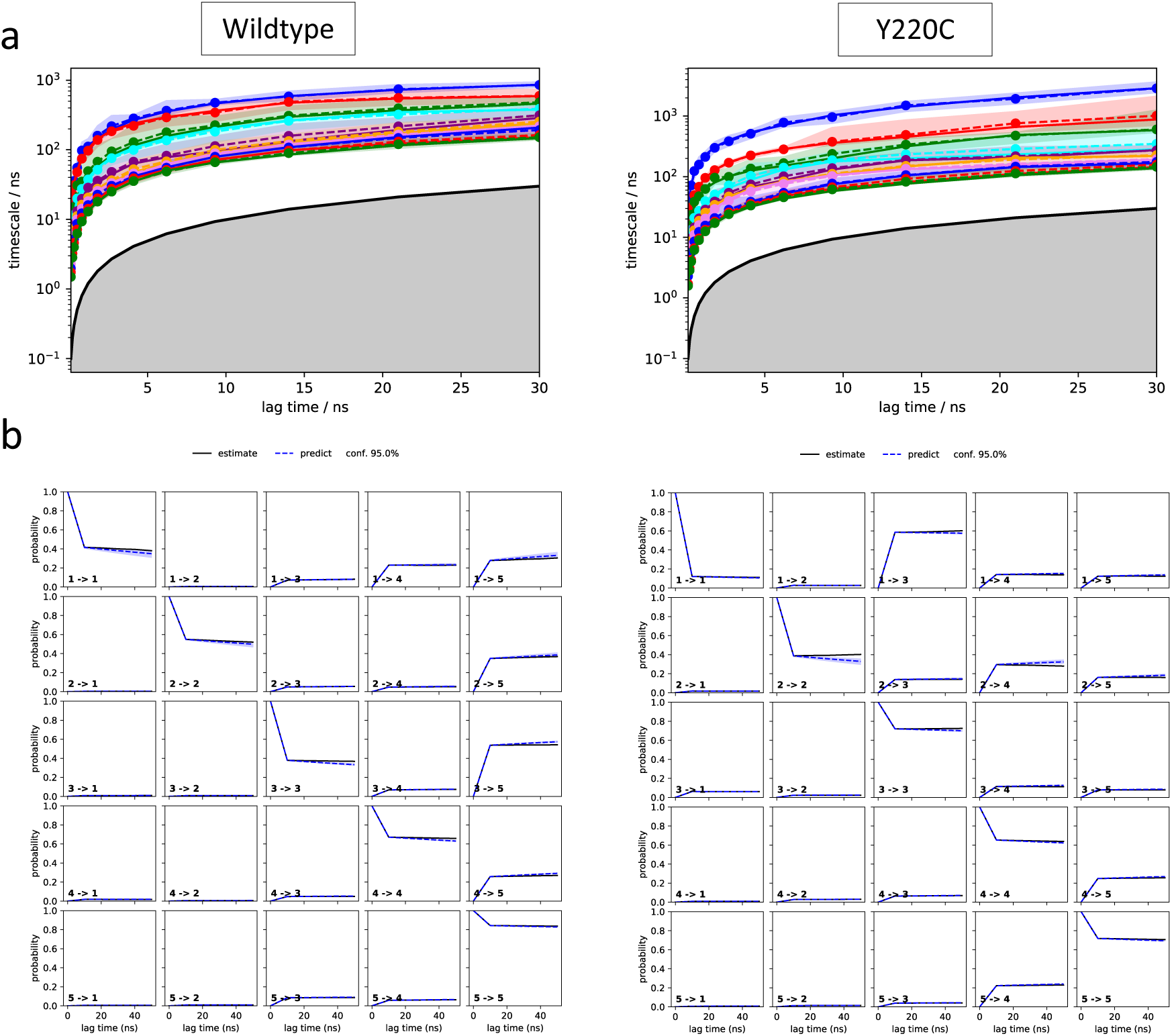
L6 MSM model validation analysis: (a) Implied timescale plots and (b) Chapman-Kolmogorov tests.

**Supplementary Figure 8.**
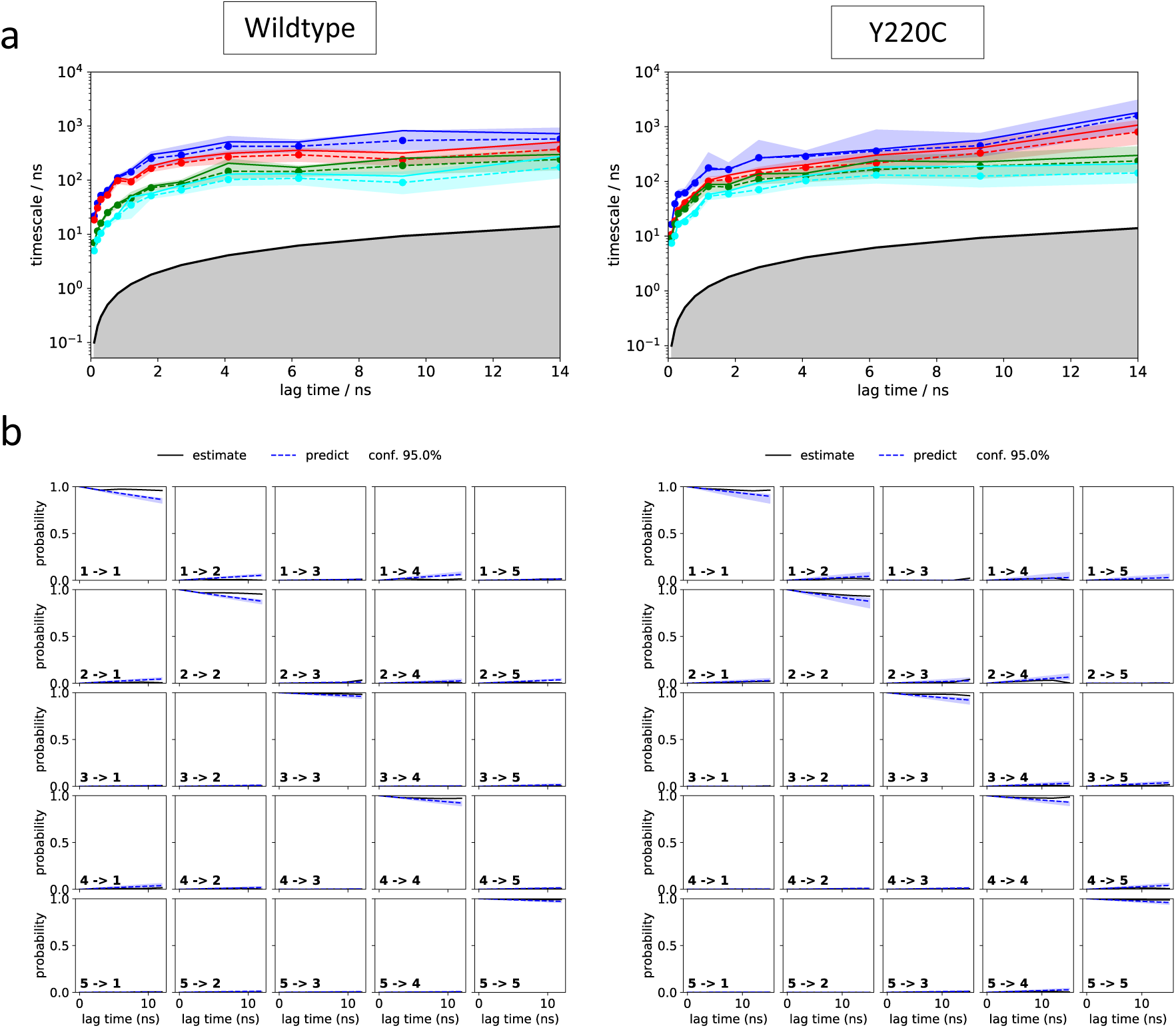
L1 HMM model validation analysis: (a) Implied timescale plots and (b) Chapman-Kolmogorov tests.

**Supplementary Figure 9.**
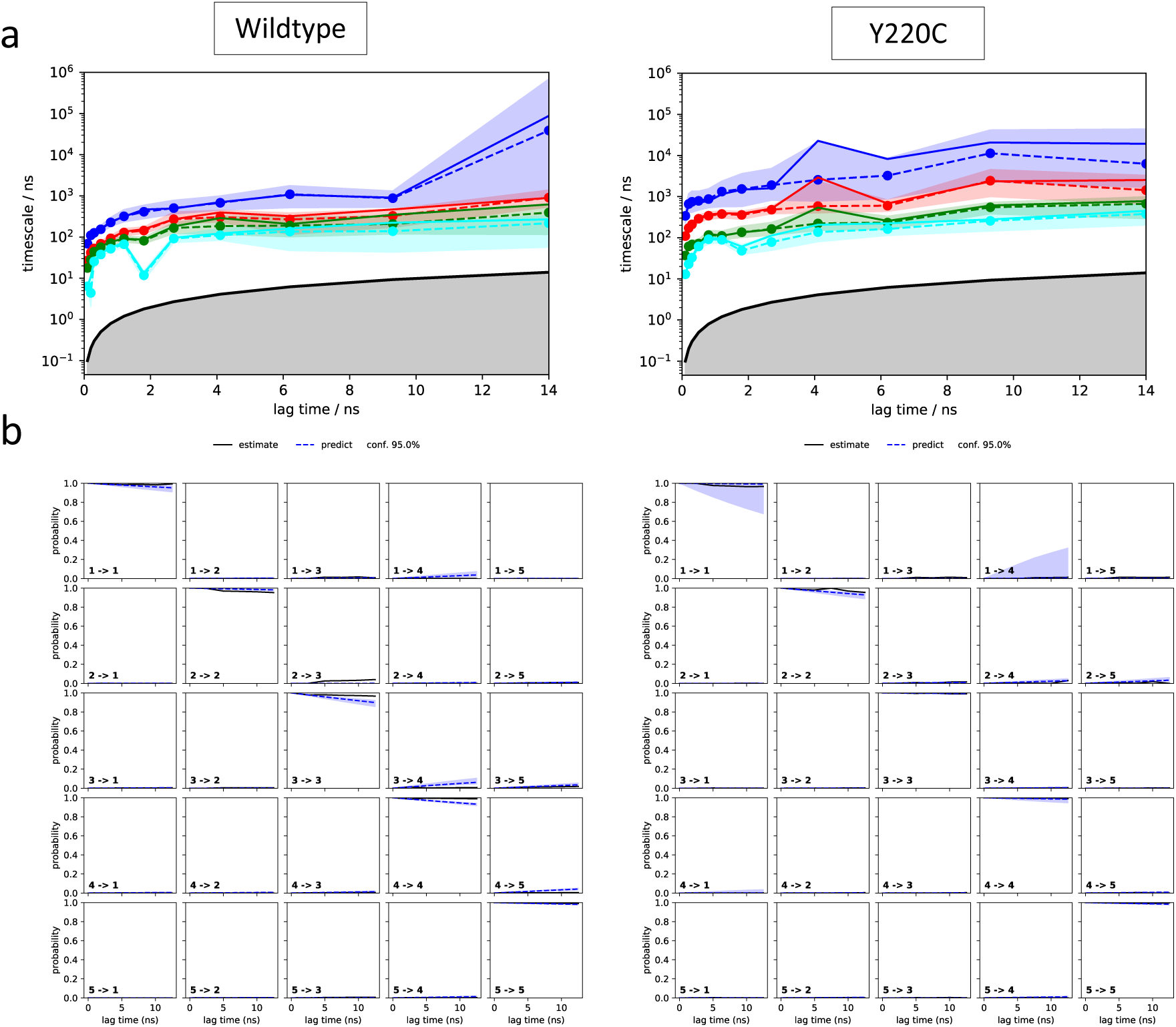
L6 HMM model validation analysis: (a) Implied timescale plots and (b) Chapman-Kolmogorov tests.

