## Supporting Information for "Markov State Models and NMR Uncover an Overlooked Allosteric Loop in p53"

$$\boldsymbol{Error\ bar\ = R2/R1 \times \sqrt{\left(\frac{R1\ error}{R1}\right)^2 + \left(\frac{R2\ error}{R2}\right)^2}}$$

NOE errors were calculated by the following equation:

$$Error\ = |NOE\ ratio| \times \sqrt{\left(\frac{NOE\ spectra\ noise}{NOE\ peak\ volume}\right)^2 + \left(\frac{reference\ spectra\ noise}{reference\ peak\ volume}\right)^2}$$

NMR generalized order parameters ($S^2$) were obtained by using *Modelfree 4.15*[16,17] in combination with *FASTModelfree.*[18] Initial estimation of *Modelfree* parameters were obtained through the programs *pdbinertia*, *r2r1_tm*, *quadric_diffusion* from the CoMD/NMR website.[19] Residues with NOE and $X^2$ values less than 0.6 and 10, respectively, were excluded from *quadric_diffusion* calculation. Our analysis was performed using the Protein Data Bank (PDB) file 2FEJ. All *FASTModelfree* analysis used an NH bond length of 1.015 Å and chemical shift entropy (CSA) value of -179 ppm similar to the previously published analysis of p53 DBD.[20] *FASTModelfree* analysis proceeded until all parameters converged.

**Supplementary Table 1.** Stepwise tICA-based selection of features for model building.

| Iteration | Number of features | Number of tICs | Correlation cutoff | Constraints for next round |
|---|---|---|---|---|
| 0 | 18,336 | - | - | Remove pairs located < 3Å or > 10 Å apart in all frames |
| 1 | 7,183 | - | - | Remove pairs with distance variance < 0.05 Å |
| 2 | 2,225 | - | - | Remove pairs involving terminal residues |
| 3 | 729 | 315 | 0.4 | None applied |
| 4 | 499 | 122 | 0.5 | None applied |
| 5 | 354 | 91 | 0.6 | None applied |
| 6 | 194 | 57 | 0.6 | None applied |
| 7 | 90 | 29 | - | Remove features that involve residues close to termini |
| 8 | 82 | 26 | - | Remove similar pairs |
| 9 | 35 | 16 | 0.75 | Remove similar pairs |
| Final | 24 | 13 | | |

**Supplementary Table 2.** Pairs used for featurization of the simulations for model construction

| Member 1 (Anchor residue) | Member 2 |
|---|---|
| Ser116 | Leu145 |
| Ser116 | Val147 |
| Ser116 | Thr150 |
| Ser116 | Tyr220 |
| Ser116 | Cys229 |
| Ser116 | Gly279 |
| Ser116 | Arg280 |
| Pro223 | Gly112 |
| Pro223 | Leu114 |
| Pro223 | Val143 |
| Pro223 | Leu145 |
| Pro223 | Thr230 |
| Glu224 | Pro153 |
| Glu224 | Gly154 |
| Glu224 | Cys229 |
| Glu224 | Ser260 |
| Glu224 | Ser261 |
| Gly226 | Thr155 |
| Gly226 | Arg156 |
| Gly226 | Pro219 |
| Gly226 | Tyr220 |
| Gly226 | Glu221 |
| Gly226 | Glu258 |
| Gly226 | Ser260 |

**Supplementary Table 3.** Wildtype X-ray and NMR structures used for comparison with simulations' conformational landscape

| PDB ID | Chain ID |
|---|---|
| 1GZH | C |
| 1KZY | A,B |
| 1TSR | A,B,C |
| 1TUP | A,B,C |
| 2AC0 | A,B,C,D |
| 2ADY | A,B |
| 2AHI | A,B,D |
| 2ATA | A,B,C,D |
| 2H1L | M,N,O,P,Q,R,S,T,U,V,W,X |
| 2OCJ | A,B,C,D |
| 2XWR | A,B |
| 2YBG | C,D |
| 3KMD | A,B,C,D |
| 3Q05 | A,B,C,D |
| 3TS8 | A,B,C,D |
| 4HJE | A,B,C,D |

**Supplementary Table 4.** Y220C X-ray and NMR structures used for comparison with simulations' conformational landscape

| **PDB ID** | **Chain ID** |
|---|---|
| 2J1X | A |
| 2VUK | A,B |
| 2X0U | A |
| 2X0V | A,B |
| 2X0W | A |
| 3ZME | A,B |
| 4AGL | A,B |
| 4AGM | A,B |
| 4AGN | A,B |
| 4AGO | A,B |
| 4AGP | A,B |
| 4AGQ | A,B |
| 5A7B | A,B |
| 5AB9 | A,B |
| 5ABA | A,B |
| 5AOI | A,B |
| 5AOJ | A,B |
| 5AOK | A,B |
| 5AOL | B |
| 5AOM | A,B |
| 5G4M | A,B |
| 5G4N | A,B |
| 5G4O | A,B |
| 6GGA | A,B |
| 6GGB | A,B |
| 6GGC | A,B |
| 6GGD | A,B |
| 6GGE | A,B |
| 6GGF | A,B |

**Supplementary Table 5.** Persistence of L6-S3/S4 hydrogen bonds (in % of frames in the simulation)

| Donor atom | Acceptor atom | Wildtype | Y220C |
|---|---|---|---|
| Thr149 - N | Gly225 - O | 0 | 0.7 |
| Thr149 - N | Asp227 – OD1 | 1.4 | 2.4 |
| Thr149 - N | Asp227 – OD2 | 1.3 | 3.19 |
| Cys219 – N | Thr154 - O | 97.0 | 85.1 |
| Ser226 – N | Thr149 – OG1 | 7.3 | 4.9 |
| Asp227 - N | Thr149 – OG1 | 0.2 | 1.9 |
| Thr149 – OG1* | Pro222 – O* | 0.09 | 9.0 |
| Thr149 – OG1 | Val224 – O | 0.02 | 1.0 |
| Thr149 – OG1 | Gly225 – O | 0.04 | 1 |
| Thr149 – OG1 | Ser226 – OG | 0.06 | 0.8 |
| Thr149 – OG1 | Ser226 – O | 1.5 | 0.9 |
| Thr149 – OG1 | Asp227 – OD1 | 3.7 | 6.6 |
| Thr149 – OG1 | Asp227 – OD2 | 3.8 | 7.3 |
| Thr149 – OG1 | Asp227 – O | 0.07 | 0.9 |
| Thr154 – OG1 | Cys219 - O | 0.2 | 5.8 |
| Ser226 - OG | Asp147 - O | 0.01 | 1.1 |
| Ser226 - OG | Thr54149 – OG1 | 0.3 | 1.0 |

* Interaction formed in the mutant-exclusive “sideways-bent” extended state

**Supplementary Table 6.** R1, R2, NOE, and $S^2$ values for Wildtype p53 DBD at 800 MHz field strength.

| Residue # | R1 ± error ($s^{-1}$) | R2 ± error ($s^{-1}$) | NOE ± error | S2 ± error |
|---|---|---|---|---|
| 99 | 1.485 ± 0.361 | 30.43 ± 0.654 | 0.795 ± 0.010 | † |
| 100 | 1.318 ± 0.284 | 28.36 ± 2.246 | 0.765 ± 0.012 | 0.791 ± 0.066 |
| 101 | 1.260 ± 0.275 | 31.81 ± 1.290 | 0.907 ± 0.006 | † |
| 102 | 1.209 ± 0.254 | 33.77 ± 4.850 | 0.657 ± 0.010 | 0.788 ± 0.089 |
| 103 | 1.087 ± 0.162 | 28.69 ± 0.811 | 0.711 ± 0.010 | 0.857 ± 0.027 |
| 104 | | | | |
| 105 | 1.383 ± 0.287 | 35.68 ± 1.806 | 0.717 ± 0.009 | 0.645 ± 0.141 |
| 106 | | | | |
| 107 | 1.255 ± 0.233 | 33.81 ± 2.730 | 0.820 ± 0.009 | † |
| 108 | 1.048 ± 0.213 | 42.69 ± 3.544 | 0.977 ± 0.010 | † |
| 109 | | | | |
| 110 | 1.056 ± 0.206 | 31.40 ± 4.243 | 0.830 ± 0.013 | 1.000 ± 0.080 |
| 111 | 0.806 ± 0.119 | 33.32 ± 1.186 | 0.745 ± 0.010 | 0.905 ± 0.047 |
| 112 | 0.773 ± 0.184 | ** | 1.000 ± 0.018 | |
| 113 | | 28.78 ± 2.079 | | |
| 114 | 0.979 ± 0.197 | 28.25 ± 1.185 | 0.775 ± 0.011 | 0.848 ± 0.036 |
| 115 | 1.718 ± 0.671 | 24.75 ± 3.405 | 0.379 ± 0.016 | 0.854 ± 0.091 |
| 116 | | | | |
| 117 | 1.561 ± 0.461 | 30.35 ± 4.726 | 0.779 ± 0.018 | † |
| 118 | 1.990 ± 0.508 | | 0.923 ± 0.016 | |
| 119 | | | | |
| 120 | | | | |
| 121 | 2.192 ± 0.573 | 29.47 ± 3.104 | 0.732 ± 0.014 | † |
| 122 | | | | |
| 123 | 1.407 ± 0.434 | 34.21 ± 3.314 | 1.000 ± 0.027 | |
| 124 | 1.172 ± 0.201 | 29.86 ± 0.850 | 0.876 ± 0.009 | 1.000 ± 0.024 |
| 125 | 0.923 ± 0.148 | 28.53 ± 0.271 | 0.854 ± 0.009 | 1.000 ± 0.008 |
| 126 | 1.355 ± 0.274 | 26.99 ± 2.243 | 0.913 ± 0.012 | 1.000 ± 0.063 |
| 127 | 0.884 ± 0.240 | 26.60 ± 3.197 | 0.866 ± 0.015 | 1.000 ± 0.080 |
| 128 | | | | |
| 129 | | | | |
| 130 | 0.933 ± 0.192 | 76.46 ± 22.161 | 0.930 ± 0.011 | † |
| 131 | | | | |
| 132 | 0.923 ± 0.160 | 34.34 ± 1.178 | 0.730 ± 0.009 | 0.858 ± 0.062 |
| 133 | 0.820 ± 0.205 | | 0.775 ± 0.021 | |
| 134 | 0.892 ± 0.114 | 30.62 ± 3.896 | 0.817 ± 0.013 | 1.000 ± 0.060 |

| | | | | |
|---|---|---|---|---|
| 135 | 1.061 ± 0.205 | 32.10 ± 2.578 | 0.784 ± 0.009 | 0.872 ± 0.071 |
| 136 | 0.953 ± 0.174 | 29.92 ± 5.053 | 0.764 ± 0.011 | 0.940 ± 0.078 |
| 137 | 0.860 ± 0.150 | 27.96 ± 1.204 | 0.743 ± 0.013 | 0.872 ± 0.038 |
| 138 | 0.690 ± 0.164 | 28.16 ± 3.482 | 0.885 ± 0.016 | 0.906 ± 0.089 |
| 139 | 1.054 ± 0.204 | 27.36 ± 1.265 | 0.820 ± 0.010 | 0.898 ± 0.033 |
| 140 | 0.996 ± 0.163 | 30.71 ± 1.633 | 0.785 ± 0.008 | 0.879 ± 0.042 |
| 141 | 0.925 ± 0.190 | 36.04 ± 2.876 | 0.907 ± 0.013 | 1.000 ± 0.054 |
| 142 | | | | |
| 143 | 0.827 ± 0.069 | 29.74 ± 3.598 | 0.642 ± 0.010 | 0.848 ± 0.031 |
| 144 | 1.371 ± 0.230 | | 0.541 ± 0.009 | 0.289 ± 0.279 |
| 145 | 0.692 ± 0.093 | 39.78 ± 3.833 | 0.855 ± 0.013 | 1.000 ± 0.060 |
| 146 | 0.878 ± 0.158 | 29.38 ± 1.939 | | |
| 147 | 0.950 ± 0.098 | 30.32 ± 1.871 | 0.711 ± 0.008 | 1.000 ± 0.038 |
| 148 | 0.893 ± 0.153 | 28.80 ± 2.982 | 1.000 ± 0.014 | † |
| 149 | 0.865 ± 0.145 | 27.82 ± 1.366 | 0.918 ± 0.007 | † |
| 150 | 0.979 ± 0.167 | 28.69 ± 0.651 | 0.832 ± 0.006 | 0.932 ± 0.018 |
| 151 | | | | |
| 152 | | | | |
| 153 | | | | |
| 154 | | | | |
| 155 | 0.870 ± 0.124 | 32.19 ± 2.627 | 0.689 ± 0.007 | 0.934 ± 0.055 |
| 156 | 0.823 ± 0.106 | 31.88 ± 0.328 | 0.830 ± 0.010 | 0.937 ± 0.011 |
| 157 | 0.774 ± 0.078 | 30.59 ± 1.577 | 0.858 ± 0.011 | 0.979 ± 0.037 |
| 158 | 1.037 ± 0.201 | 36.18 ± 5.200 | 0.896 ± 0.011 | 1.000 ± 0.083 |
| 159 | 1.009 ± 0.270 | 33.48 ± 3.200 | 0.833 ± 0.013 | 1.000 ± 0.053 |
| 160 | 1.040 ± 0.178 | 33.32 ± 1.551 | 0.983 ± 0.014 | † |
| 161 | 0.759 ± 0.168 | ** | 0.609 ± 0.016 | |
| 162 | 0.887 ± 0.173 | ** | 0.864 ± 0.011 | |
| 163 | 1.114 ± 0.299 | * | 0.978 ± 0.023 | |
| 164 | | | | |
| 165 | | | | |
| 166 | | | | |
| 167 | | | | |
| 168 | | 32.56 ± 0.841 | | |
| 169 | 1.164 ± 0.257 | 38.47 ± 2.541 | 0.849 ± 0.010 | 1.000 ± 0.046 |
| 170 | 1.885 ± 0.445 | | 0.645 ± 0.010 | |
| 171 | 1.265 ± 0.262 | 35.89 ± 2.385 | 0.836 ± 0.007 | 1.000 ± 0.054 |
| 172 | 1.123 ± 0.206 | 31.12 ± 2.398 | 0.835 ± 0.010 | 1.000 ± 0.051 |
| 173 | 1.220 ± 0.170 | * | 0.871 ± 0.019 | |
| 174 | 1.208 ± 0.277 | 37.97 ± 4.934 | 1.000 ± 0.011 | † |

| | | | | |
|---|---|---|---|---|
| 175 | 1.256 ± 0.365 | ** | 0.638 ± 0.016 | |
| 176 | | | | |
| 177 | | | | |
| 178 | | 30.25 ± 0.508 | 1.000 ± 0.006 | |
| 179 | 0.845 ± 0.094 | 34.63 ± 4.812 | 0.871 ± 0.011 | 1.000 ± 0.061 |
| 180 | 1.100 ± 0.195 | 37.32 ± 0.588 | 0.855 ± 0.010 | 1.000 ± 0.014 |
| 181 | 0.984 ± 0.166 | 38.27 ± 1.342 | 0.888 ± 0.010 | 1.000 ± 0.022 |
| 182 | 1.325 ± 0.231 | 33.77 ± 0.584 | 0.826 ± 0.008 | † |
| 183 | | | | |
| 184 | 1.862 ± 0.468 | | 0.855 ± 0.011 | |
| 185 | 0.959 ± 0.191 | 30.68 ± 1.257 | 0.824 ± 0.008 | 0.938 ± 0.029 |
| 186 | 1.831 ± 0.361 | 25.83 ± 0.958 | 0.876 ± 0.009 | 0.776 ± 0.027 |
| 187 | 1.544 ± 0.275 | 27.05 ± 1.468 | 0.669 ± 0.008 | 0.540 ± 0.131 |
| 188 | 1.250 ± 0.223 | 25.80 ± 1.758 | 0.544 ± 0.007 | 0.641 ± 0.118 |
| 189 | 1.000 ± 0.172 | 25.45 ± 2.785 | 0.944 ± 0.017 | † |
| 190 | | | | |
| 191 | | | | |
| 192 | 1.266 ± 0.273 | 21.52 ± 2.605 | 0.776 ± 0.011 | 0.675 ± 0.071 |
| 193 | 1.133 ± 0.278 | 36.58 ± 1.683 | 0.837 ± 0.011 | 1.000 ± 0.028 |
| 194 | | | | |
| 195 | 1.669 ± 0.747 | | 0.815 ± 0.022 | |
| 196 | 0.821 ± 0.150 | | 0.719 ± 0.013 | |
| 197 | | | | |
| 198 | 0.830 ± 0.116 | 28.91 ± 1.925 | 0.725 ± 0.010 | 0.904 ± 0.042 |
| 199 | 1.069 ± 0.193 | 26.88 ± 0.249 | 0.782 ± 0.010 | 0.889 ± 0.010 |
| 200 | 1.137 ± 0.195 | 26.18 ± 0.654 | 0.779 ± 0.007 | 0.798 ± 0.069 |
| 201 | 1.060 ± 0.314 | 29.00 ± 1.992 | 0.807 ± 0.019 | 0.851 ± 0.053 |
| 202 | 0.799 ± 0.098 | 31.51 ± 1.085 | 0.836 ± 0.008 | 1.000 ± 0.021 |
| 203 | 0.844 ± 0.088 | 28.17 ± 1.721 | 0.653 ± 0.007 | 0.837 ± 0.033 |
| 204 | 0.839 ± 0.057 | 31.78 ± 1.478 | 0.795 ± 0.010 | 0.906 ± 0.024 |
| 205 | 0.755 ± 0.105 | 33.81 ± 1.180 | 0.890 ± 0.010 | 1.000 ± 0.021 |
| 206 | 0.968 ± 0.169 | 33.20 ± 3.624 | 0.656 ± 0.012 | 0.815 ± 0.072 |
| 207 | | | | |
| 208 | 1.005 ± 0.186 | 20.76 ± 1.623 | 0.636 ± 0.008 | 0.761 ± 0.058 |
| 209 | 0.955 ± 0.213 | 30.69 ± 2.027 | | |
| 210 | 1.116 ± 0.156 | 28.50 ± 1.283 | 0.692 ± 0.007 | 0.824 ± 0.072 |
| 211 | 0.870 ± 0.173 | 27.78 ± 1.765 | 0.756 ± 0.015 | 0.911 ± 0.044 |
| 212 | 0.986 ± 0.130 | 33.73 ± 4.092 | 0.807 ± 0.012 | 0.862 ± 0.042 |
| 213 | 1.062 ± 0.255 | 32.79 ± 0.978 | | |
| 214 | 1.530 ± 0.371 | 27.03 ± 7.639 | 0.830 ± 0.017 | 1.000 ± 0.127 |

| | | | | |
|---|---|---|---|---|
| 215 | 1.065 ± 0.218 | * | 0.802 ± 0.014 | |
| 216 | 1.323 ± 0.276 | 30.35 ± 1.261 | 0.761 ± 0.006 | 0.659 ± 0.088 |
| 217 | 0.824 ± 0.166 | 29.17 ± 5.349 | 1.000 ± 0.012 | † |
| 218 | 0.869 ± 0.111 | 35.61 ± 4.136 | 0.844 ± 0.011 | 1.000 ± 0.064 |
| 219 | | | | |
| 220 | 0.778 ± 0.090 | 25.48 ± 2.771 | 0.919 ± 0.010 | † |
| 221 | 0.694 ± 0.066 | 34.36 ± 2.672 | 0.766 ± 0.008 | 0.948 ± 0.025 |
| 222 | | | | |
| 223 | | | | |
| 224 | 0.982 ± 0.124 | 24.56 ± 0.762 | 0.482 ± 0.006 | 0.790 ± 0.026 |
| 225 | | | | |
| 226 | 1.173 ± 0.222 | 25.92 ± 0.488 | 0.303 ± 0.009 | 0.755 ± 0.017 |
| 227 | | | | |
| 228 | 1.476 ± 0.368 | 24.11 ± 2.166 | 0.528 ± 0.009 | 0.664 ± 0.065 |
| 229 | 1.143 ± 0.219 | 30.11 ± 1.104 | 0.805 ± 0.006 | 0.867 ± 0.025 |
| 230 | 1.375 ± 0.193 | 16.88 ± 0.410 | 0.727 ± 0.007 | 0.452 ± 0.017 |
| 231 | 0.909 ± 0.143 | 26.67 ± 0.665 | 0.891 ± 0.009 | 0.962 ± 0.022 |
| 232 | 0.773 ± 0.092 | 31.50 ± 1.835 | | |
| 233 | 1.212 ± 0.167 | 13.47 ± 0.814 | 0.570 ± 0.007 | 0.360 ± 0.026 |
| 234 | 0.870 ± 0.164 | 26.40 ± 3.284 | 0.813 ± 0.013 | 0.909 ± 0.052 |
| 235 | 0.804 ± 0.132 | 32.41 ± 2.290 | 0.769 ± 0.010 | 0.933 ± 0.041 |
| 236 | 0.889 ± 0.224 | 27.46 ± 2.910 | 0.667 ± 0.015 | 0.889 ± 0.066 |
| 237 | 0.993 ± 0.222 | ** | 0.789 ± 0.019 | |
| 238 | 0.922 ± 0.149 | 38.71 ± 2.767 | 0.828 ± 0.008 | 0.966 ± 0.037 |
| 239 | 0.910 ± 0.196 | * | 0.821 ± 0.020 | |
| 240 | 0.817 ± 0.265 | 30.28 ± 4.720 | 1.000 ± 0.020 | † |
| 241 | 1.390 ± 0.390 | 37.53 ± 1.806 | 0.818 ± 0.014 | 1.000 ± 0.053 |
| 242 | 1.290 ± 0.237 | 35.10 ± 1.120 | 0.857 ± 0.008 | 1.000 ± 0.028 |
| 243 | | ** | 0.641 ± 0.009 | |
| 244 | 0.691 ± 0.349 | | 0.441 ± 0.024 | |
| 245 | 1.356 ± 0.316 | 29.32 ± 4.274 | 0.752 ± 0.011 | 0.750 ± 0.098 |
| 246 | | | | |
| 247 | | | | |
| 248 | | | | |
| 249 | | | | |
| 250 | | | | |
| 251 | 1.238 ± 0.228 | ** | 0.891 ± 0.017 | |
| 252 | 0.877 ± 0.152 | 32.91 ± 3.637 | 1.000 ± 0.016 | † |
| 253 | 0.777 ± 0.093 | 30.40 ± 3.035 | 0.794 ± 0.011 | 0.952 ± 0.041 |
| 254 | 0.905 ± 0.152 | 27.89 ± 6.521 | 0.749 ± 0.010 | 0.927 ± 0.060 |

| | | | | |
|---|---|---|---|---|
| 255 | 0.735 ± 0.142 | 35.66 ± 3.189 | 0.804 ± 0.011 | 0.953 ± 0.045 |
| 256 | 1.019 ± 0.164 | ** | 0.880 ± 0.011 | |
| 257 | 0.821 ± 0.151 | 32.81 ± 0.732 | 0.978 ± 0.014 | † |
| 258 | 0.797 ± 0.097 | 30.05 ± 4.498 | 0.744 ± 0.008 | 0.921 ± 0.038 |
| 259 | | | | |
| 260 | 1.313 ± 0.234 | 28.84 ± 1.114 | 0.708 ± 0.009 | 0.720 ± 0.116 |
| 261 | 1.147 ± 0.195 | 30.43 ± 0.760 | 0.679 ± 0.006 | 0.805 ± 0.096 |
| 262 | 1.112 ± 0.200 | 28.30 ± 1.457 | 0.684 ± 0.007 | 0.766 ± 0.091 |
| 263 | 0.873 ± 0.088 | 29.22 ± 1.271 | 0.763 ± 0.006 | 0.884 ± 0.031 |
| 264 | 0.935 ± 0.140 | 25.10 ± 1.018 | 0.836 ± 0.008 | 0.939 ± 0.024 |
| 265 | 0.870 ± 0.112 | 24.07 ± 2.226 | 0.679 ± 0.011 | 0.843 ± 0.041 |
| 266 | 0.767 ± 0.099 | 35.92 ± 4.275 | 0.798 ± 0.012 | 0.979 ± 0.050 |
| 267 | 1.042 ± 0.181 | 34.64 ± 3.027 | 0.706 ± 0.013 | 0.777 ± 0.079 |
| 268 | 0.755 ± 0.134 | 36.05 ± 0.948 | 0.777 ± 0.009 | 0.913 ± 0.051 |
| 269 | 0.949 ± 0.176 | 28.42 ± 1.107 | 0.839 ± 0.008 | 0.941 ± 0.024 |
| 270 | 0.887 ± 0.130 | 32.60 ± 2.098 | 0.834 ± 0.010 | 1.000 ± 0.036 |
| 271 | 0.876 ± 0.163 | 23.84 ± 2.207 | 0.990 ± 0.011 | † |
| 272 | | | | |
| 273 | 0.907 ± 0.154 | 33.35 ± 4.561 | 0.739 ± 0.012 | 0.948 ± 0.071 |
| 274 | 1.052 ± 0.308 | ** | 0.748 ± 0.012 | |
| 275 | 0.917 ± 0.159 | 32.24 ± 3.800 | 0.831 ± 0.011 | 1.000 ± 0.068 |
| 276 | | | | |
| 277 | 1.334 ± 0.288 | 30.05 ± 1.155 | 0.700 ± 0.007 | 0.872 ± 0.038 |
| 278 | | | | |
| 279 | 1.037 ± 0.172 | 37.30 ± 4.563 | 0.704 ± 0.010 | 0.775 ± 0.079 |
| 280 | 0.969 ± 0.190 | 31.54 ± 2.781 | 0.885 ± 0.011 | 1.000 ± 0.054 |
| 281 | 1.014 ± 0.149 | 31.55 ± 1.412 | 0.893 ± 0.009 | 0.995 ± 0.028 |
| 282 | 0.907 ± 0.142 | 35.02 ± 4.009 | 1.000 ± 0.012 | † |
| 283 | 0.810 ± 0.133 | 32.90 ± 0.830 | 0.730 ± 0.007 | 0.892 ± 0.053 |
| 284 | 1.009 ± 0.151 | 30.11 ± 0.433 | 0.635 ± 0.005 | 0.899 ± 0.013 |
| 285 | 0.930 ± 0.124 | 29.24 ± 1.380 | 0.749 ± 0.007 | 0.879 ± 0.039 |
| 286 | 0.838 ± 0.105 | 32.58 ± 2.887 | 0.821 ± 0.009 | 0.909 ± 0.027 |
| 287 | 0.845 ± 0.107 | 28.96 ± 1.542 | 0.694 ± 0.007 | 0.904 ± 0.037 |
| 288 | 0.886 ± 0.101 | 32.04 ± 0.495 | 0.764 ± 0.006 | 0.853 ± 0.044 |
| 289 | 0.844 ± 0.066 | 27.71 ± 0.712 | 0.759 ± 0.006 | 0.868 ± 0.018 |
| 290 | | | | |
| 291 | | | | |
| 292 | 1.935 ± 0.323 | 18.56 ± 0.266 | 0.518 ± 0.004 | |
| 293 | 2.608 ± 0.507 | 16.17 ± 0.553 | 0.382 ± 0.005 | |
| 294 | 2.297 ± 0.365 | 12.48 ± 0.251 | 0.489 ± 0.002 | |

| | | | |
|---|---|---|---|
| 295 | | | |
| 296 | | | |
| 297 | | | |
| 298 | 3.202 ± 0.661 | 9.95 ± 0.387 | 0.380 ± 0.003 |
| 299 | 2.166 ± 0.255 | 6.84 ± 0.121 | 0.322 ± 0.002 |
| 300 | | | |
| 301 | | | |
| 302 | | | |
| 303 | 3.522 ± 0.807 | 9.57 ± 0.600 | 0.374 ± 0.009 |
| 304 | | | |
| 305 | 3.965 ± 0.757 | 10.16 ± 0.646 | 0.281 ± 0.005 |
| 306 | 4.271 ± 0.877 | 8.21 ± 0.434 | 0.286 ± 0.005 |
| 307 | 3.801 ± 0.700 | 8.44 ± 0.224 | 0.256 ± 0.004 |
| 308 | 2.345 ± 0.363 | 5.96 ± 0.151 | 0.012 ± 0.002 |
| 309 | | | |
| 310 | 3.730 ± 0.835 | 7.93 ± 0.599 | |
| 311 | | 10.82 ± 0.652 | 0.291 ± 0.005 |

* Signal present up to .030 seconds
** Signal present up to .050 seconds
† *Modelfree* calculation did not assign a model to residue.
Blank spaces correspond to residues with relaxation values that could not be obtained.

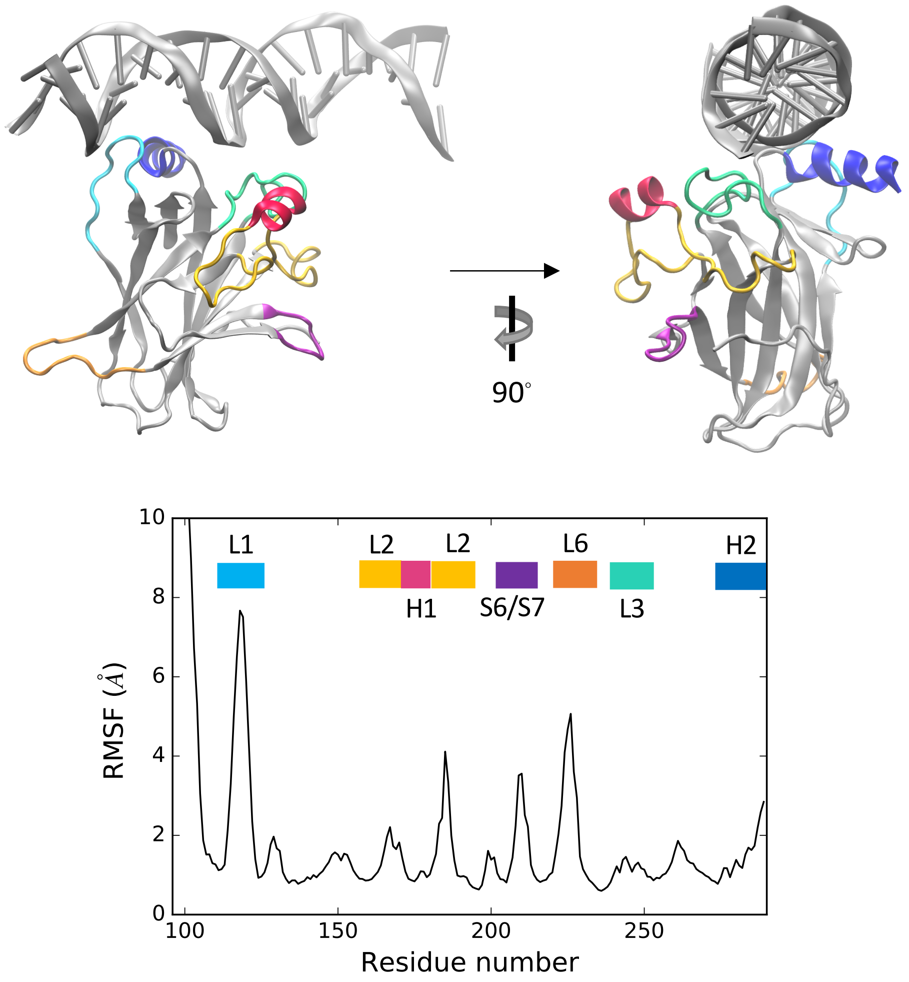


**Supplementary Figure 1.** Alpha carbon RMSF. Functionally important structural motifs are highlighted in the DNA-bound structure (top panel).

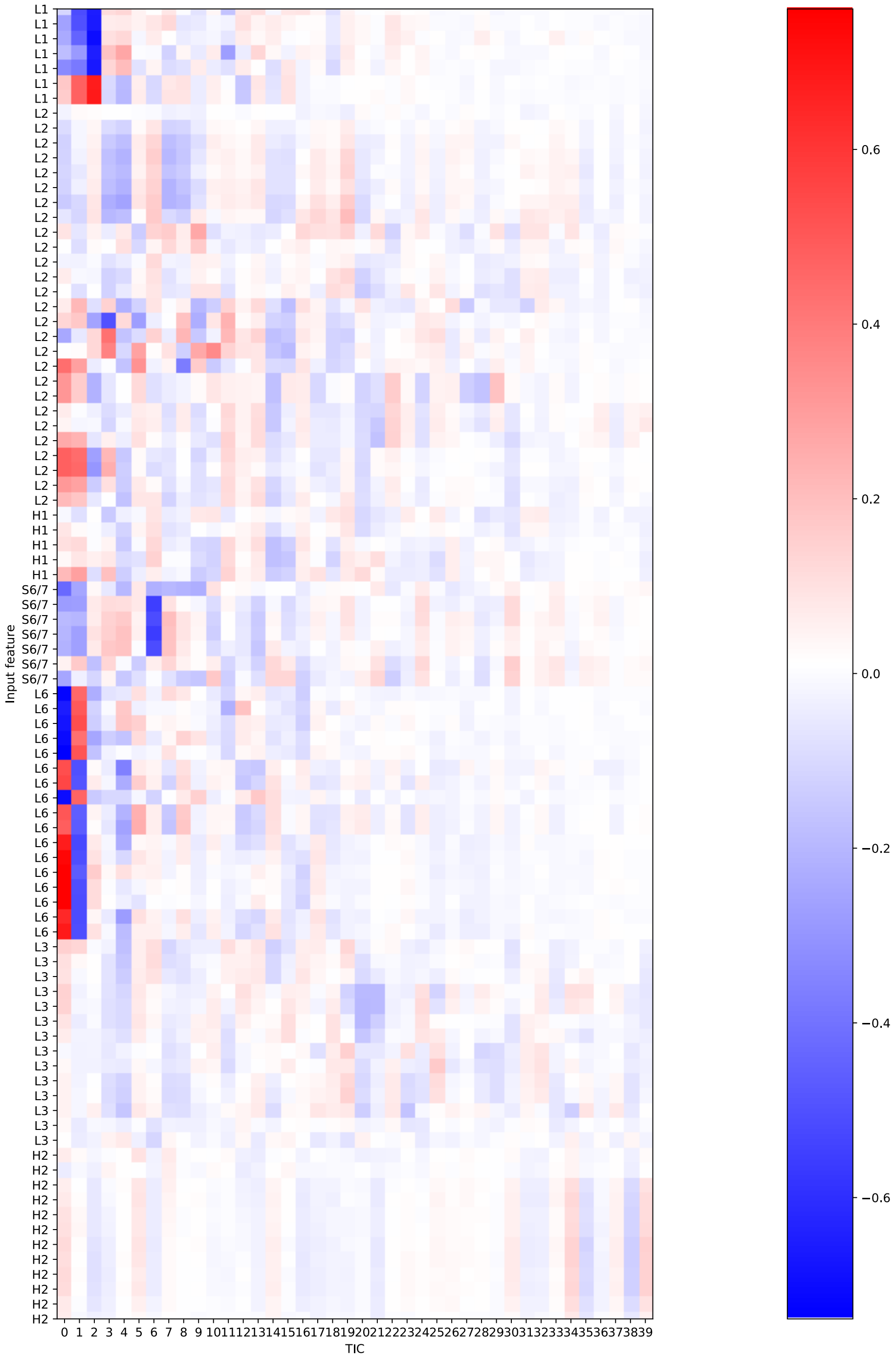


**Supplementary Figure 2.** tICA correlation for features incorporating functionally-important motifs in the protein (H1, H2, L2, L3, S6/7) in addition to L1 and L6.

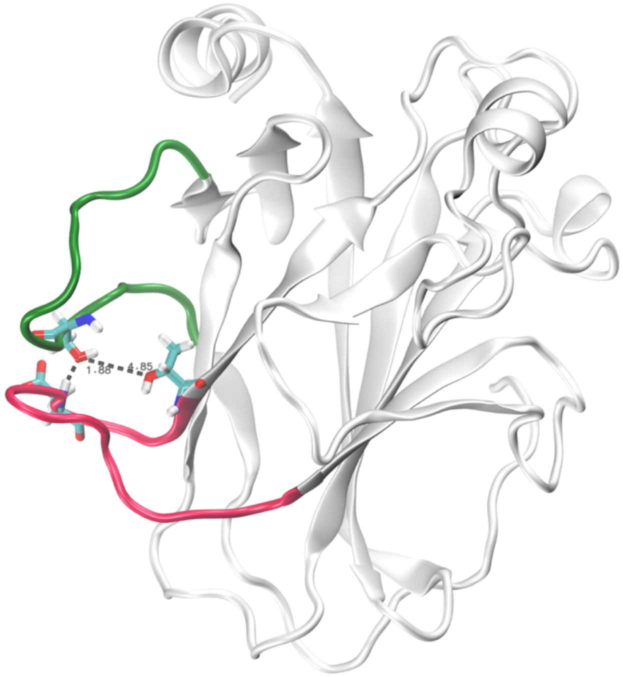


**Supplementary Figure 3.** Example of frame exhibiting most stable intra-loop hydrogen bonds, involving Ser116 in L1 and Asp228 or Thr231 in L6.

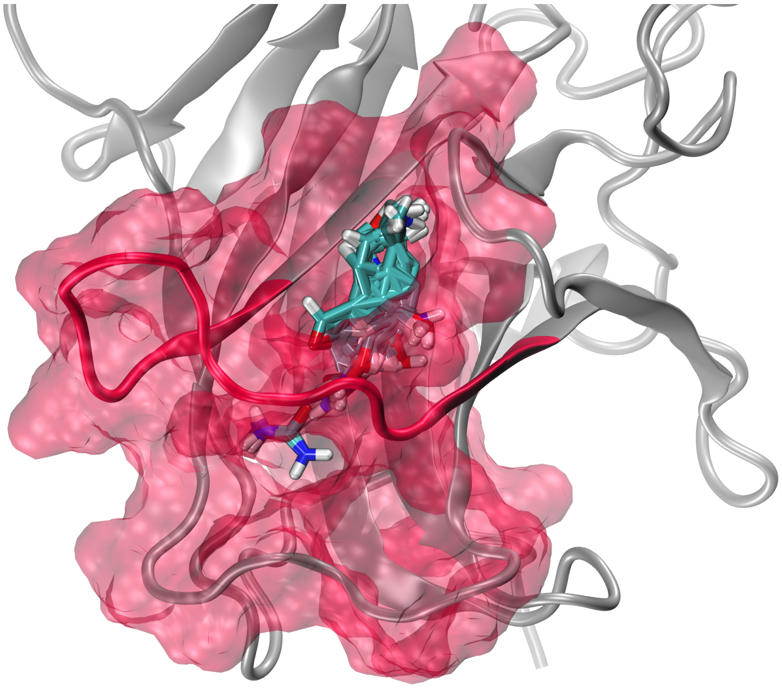

**Supplementary Figure 4.** Representation of the cryptic channel spanning loop L6 in the recessed Y220C metastable state. FTMap[11] probes indicating hotspots for drug binding are shown in licorice.

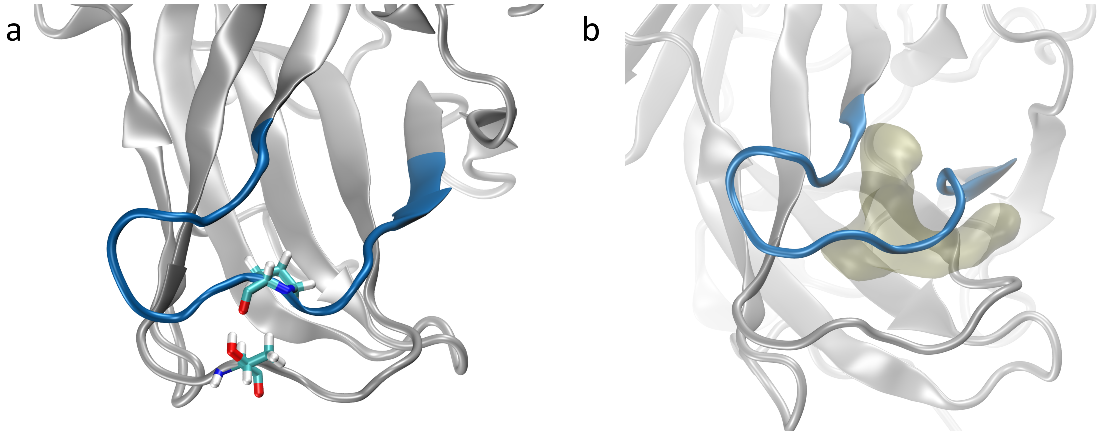


**Supplementary Figure 5.** (a) Representation of the Thr149-Pro222 interaction thought to stabilize the bent L6 conformation observed in the mutant-exclusive states. (b) Surface representation of the L6 pocket in the mutant-exclusive states.

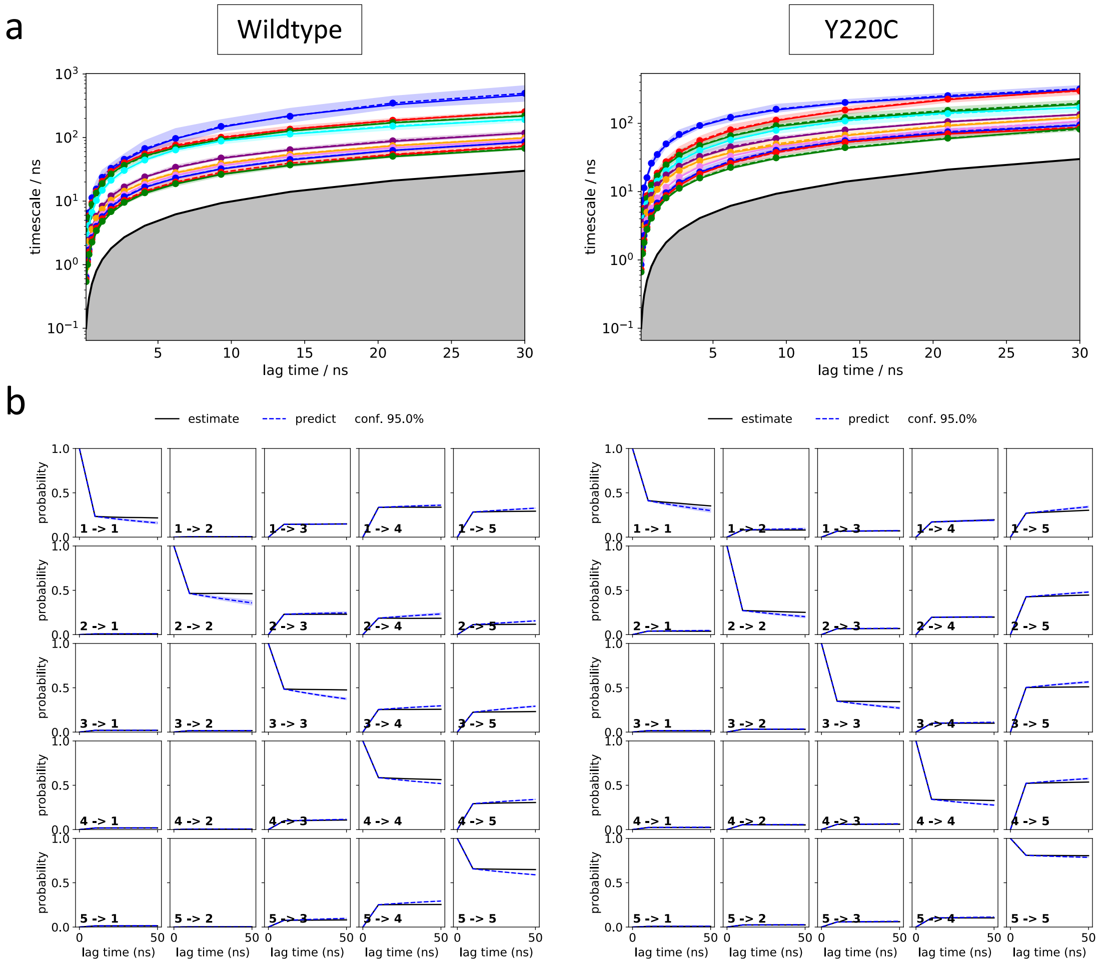


**Supplementary Figure 6.** L1 MSM model validation analysis: (a) Implied timescale plots and (b) Chapman-Kolmogorov tests.

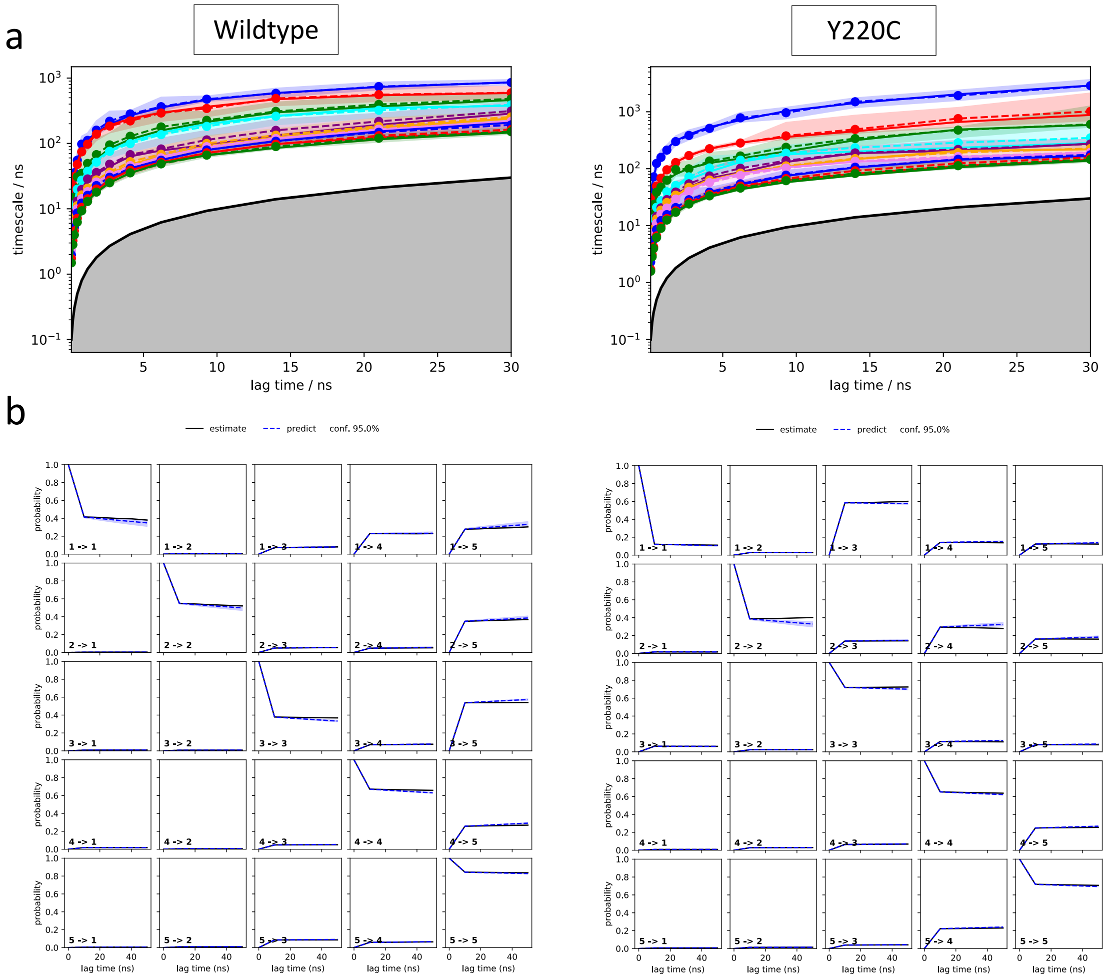


**Supplementary Figure 7.** L6 MSM model validation analysis: (a) Implied timescale plots and (b) Chapman-Kolmogorov tests.

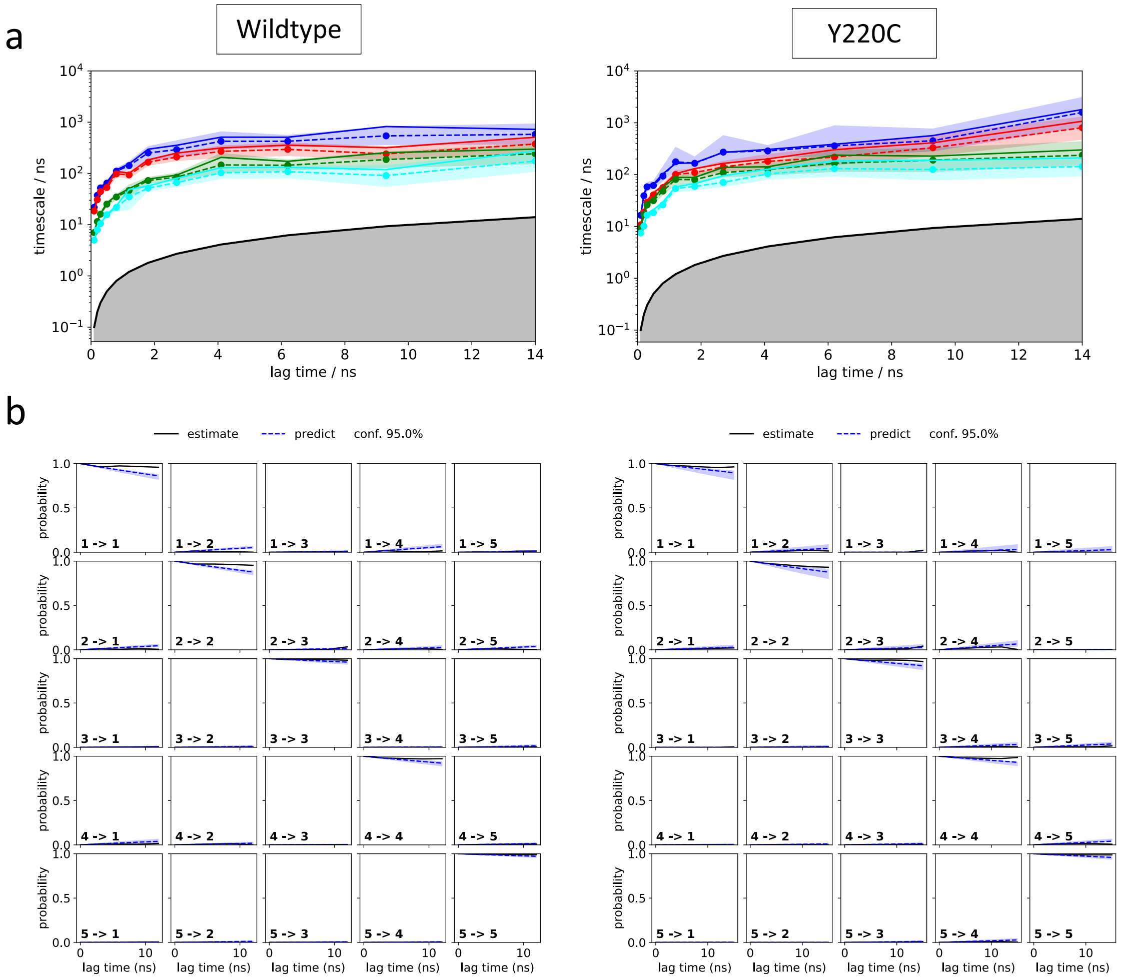


**Supplementary Figure 8.** L1 HMM model validation analysis: (a) Implied timescale plots and (b) Chapman-Kolmogorov tests.

a

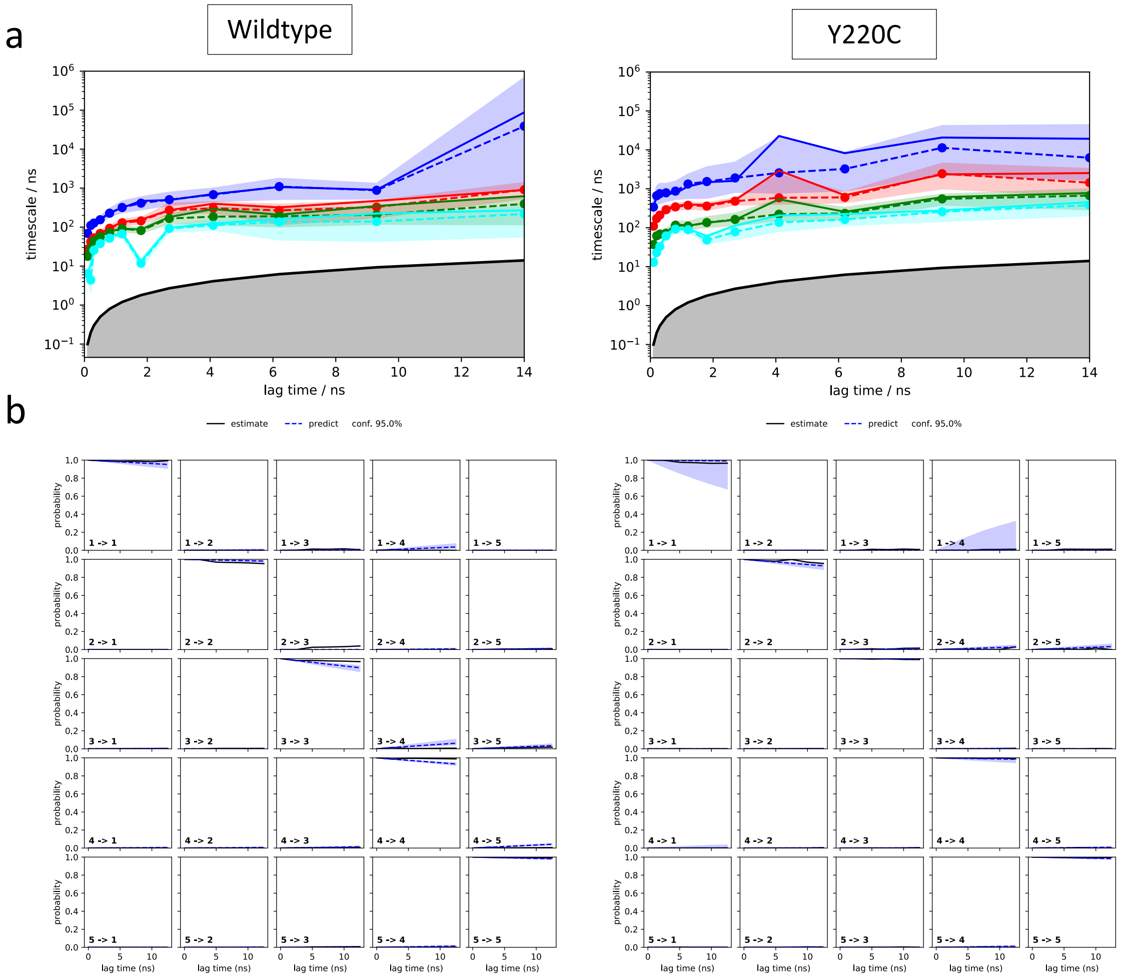


**Supplementary Figure 9.** L6 HMM model validation analysis: (a) Implied timescale plots and (b) Chapman-Kolmogorov tests.
